# Kinome focused CRISPR-Cas9 screens in African ancestry patient-derived breast cancer organoids identifies essential kinases and synergy of EGFR and FGFR1 inhibition

**DOI:** 10.1101/2023.12.11.570465

**Authors:** Florencia P. Madorsky Rowdo, Rachel Martini, Sarah Ackermann, Colin Tang, Marvel Tranquille, Adriana Irizarry, Ilkay Us, Omar Alawa, Jenna Moyer, Michael Sigouros, John Nguyen, Majd Al Assaad, Esther Cheng, Paula S. Ginter, Jyothi Manohar, Brian Stonaker, Richard Boateng, Joseph K. Oppong, Ernest K. Adjei, Baffour Awuah, Ishmael Kyei, Frances S. Aitpillah, Michael O. Adinku, Kwasi Ankomah, Ernest B. Osei-Bonsu, Kofi K. Gyan, Syed Hoda, Lisa Newman, Juan Miguel Mosquera, Andrea Sboner, Olivier Elemento, Lukas E. Dow, Melissa B. Davis, M. Laura Martin

## Abstract

Precision medicine approaches to cancer treatment aim to exploit genomic alterations that are specific to individual patients to tailor therapy strategies. These alterations are usually revealed via next generation sequencing of the tumor tissue. Yet, it is clear that some targetable genes and pathways are essential for tumor cell viability even in the absence of direct genomic alterations. This is especially important in under-represented populations, whose mutational landscape and determinants of response to existing therapies are poorly characterized due to limited inclusion in clinical trials and studies. One way to reveal tumor essential genes is with genetic screens. Most screens are conducted on cell lines that bear little resemblance to patient tumors, after years of culture in non-physiological conditions. To address this problem, we aimed to develop a CRISPR screening pipeline in 3D-grown patient-derived tumor organoid (PDTO) models. We focused on identifying essential kinases that may translate to options for targeted therapies, including combination therapies. We first established a breast cancer PDTO biobank focused on underrepresented populations, including West African patients. We then performed a negative selection kinome-focused CRISPR screen to identify kinases essential for organoid growth and potential targets for combination therapy with EGFR or MEK inhibitors. We identified several previously unidentified kinase targets and showed that combination of FGFR1 and EGFR inhibitors synergizes to block organoids proliferation. Together these data demonstrate feasibility of CRISPR-based genetic screens in patient-derived tumor models, including PDTOs from under-represented cancer patients, and identify new targets for cancer therapy.

## Introduction

Precision oncology is primarily based on the idea that genetic alterations identified in patient’s tumors can be targeted with drugs matching those alterations. This paradigm has generated several successful therapeutics, including targeting *BCR::ABL* fusions in chronic myeloid leukemia, BRAF^V600E^ in melanoma, mutated EGFR in lung cancer patients and many more. Nevertheless, data from several precision oncology initiatives [1–3] have revealed that a substantial number of patients do not experience advantages even when actionable alterations are present, as malignant processes can be driven by non-genetic mechanisms. Functional precision medicine is emerging as a complementary strategy to connect genotype and phenotype by creating in vitro models of individual tumors and directly expose them to drugs [4].

For functional precision medicine to work and reliably complement genomic testing, there is a need for personalized models that accurately represent the characteristics of individual tumors [5]. One promising approach is the use of patient-derived tumor organoids (PDTOs). PDTOs are three-dimensional cellular models grown in culture *ex vivo* from tumor tissue obtained directly from patients. This approach retains many cell–cell and cell–matrix interactions mimicking the original tumor better than cells grown in monolayers. PDTOs have been established for different cancer types, including breast cancer [6, 7]. In recent years, organoid technology has been leveraged to test the efficacy of antitumoral drugs, even allowing for combinatorial drug screens. Importantly, several studies have shown that PDTO drug responses correlate with patient responses [8].

Although largely unexplored, we argue that functional precision medicine approaches are particularly relevant in under-represented populations, whose mutational landscape and determinants of response to existing therapies are poorly characterized due to limited inclusion in clinical trials and studies. To address this gap and establish a new paradigm for such underserved populations, we first established a breast cancer PDTO biobank focused on African American and African patients. We chose breast cancer as the focus of the biobank because epidemiological studies of breast cancer incidence and prevalence across the global African Diaspora have shown that women of African descent have worse clinical outcomes compared to women of European descent [9–13]. Factors that likely contribute to AA women having a disproportionately higher breast cancer mortality rate include tumor characteristics [14] such as the higher rate of Triple Negative Breast cancer (TNBC) and earlier onset with later stages compared to other patient groups [15–17]. For our PDTO biobank, African breast cancer tumor samples were collected as part of an international collaboration with the International Center for the Study of Breast Cancer Subtypes (ICSBCS) [18].

Functional genomics screening represents a potent method for systematically investigating the roles of genes and their functions, and its implementation has been notably simplified by the emergence of CRISPR-based technologies. Genome-wide CRISPR screening in human cancer cell lines using pooled lentiviral libraries have been used extensively [19, 20], however, its application to human organoids has been technically challenging due to several factors including low growth rate of 3D cultures. Several studies have shown that targeted CRISPR mutagenesis can be successfully performed in human PDTOs [21–23]. The use of CRISPR-Cas9 to target known functional domains of candidate genes yielded successful loss-of-function (LOF) mutations to establish the roles these genes play in cell viability, proliferation, and response to drug treatments [24, 25]. To maximize the efficiency of gene disruption, we use a library of gRNAs targeting known functional domains in kinases [25–27].

We chose to focus our screen on kinase genes because (1) these are actionable targets, with numerous approved and investigational compounds targeting many kinases and (2) in a recent transcriptomic analysis of TNBC tumors among women of African ancestry, a large number of kinases were found to be upregulated compared to tumors from European Americans [28, 29]. We reasoned that biological differences in breast tumors from patients with West African genetic ancestry could be exploited to identify novel therapeutic targets that may specifically benefit patients of West African descent, even across the entire African diaspora. Thus, in this study we assessed which kinases are essential for tumor viability or directly impact the response to targeted therapies targeting frequently activated signaling pathways in PDTOs derived from West African breast cancer patients.

## Results

### Establishing an African ancestry breast cancer PDTO biobank

We first established a breast cancer PDTO biobank from an underserved population of patients, including African American (AA) and African (Ghanaian) patients. Seventeen (17) Ghanaian PDTOs cultures were established following collections through the ICSBCS [18, 30], resulting in 8 short-term and 5 long-term (13 total) cultures, with a success rate of 76%. In addition, we established 69 breast cancer PDTOs from samples collected at Weill Cornell Medicine (WCM), including 8 long-term cultures from African American patients. **Supplementary Table 1** describes the characteristics of the patient and PDTO lines in this biobank.

For all PDTOs, we performed histopathological review of hematoxylin and eosin (H&E) stained sections to determine if tumor morphology and cellular characteristics were maintained in the organoid. Immunohistochemistry (IHC) staining for Estrogen receptor (ER), Progesterone receptor (PR) and HER2 in the tumor tissue and matched PDTOs was performed to determine whether each culture reflected the molecular profile of the original tumor. Representative images of the 2 Ghanaian PDTOs used in subsequent experiments, ICSBCS002 and ICSBCS007, are shown in **Figure 1A**. ICSBCS007 PDTO matched the tumor histopathological characteristics, being ER+PR-HER2-. In contrast, ICSBCS002 demonstrates a discrepant profile, with tumor being ER positive moderate to strong in 95% of the cells, and PR positive in 1-5% of cells, while the corresponding PDTO demonstrate ER and PR negative staining. HER2 is consistently negative in both the PDTO and matching tumor. All other breast PDTO lines used in this study showed morphological consistency with original tumors (**Supplementary Figure 1A**).

**Figure 1.**
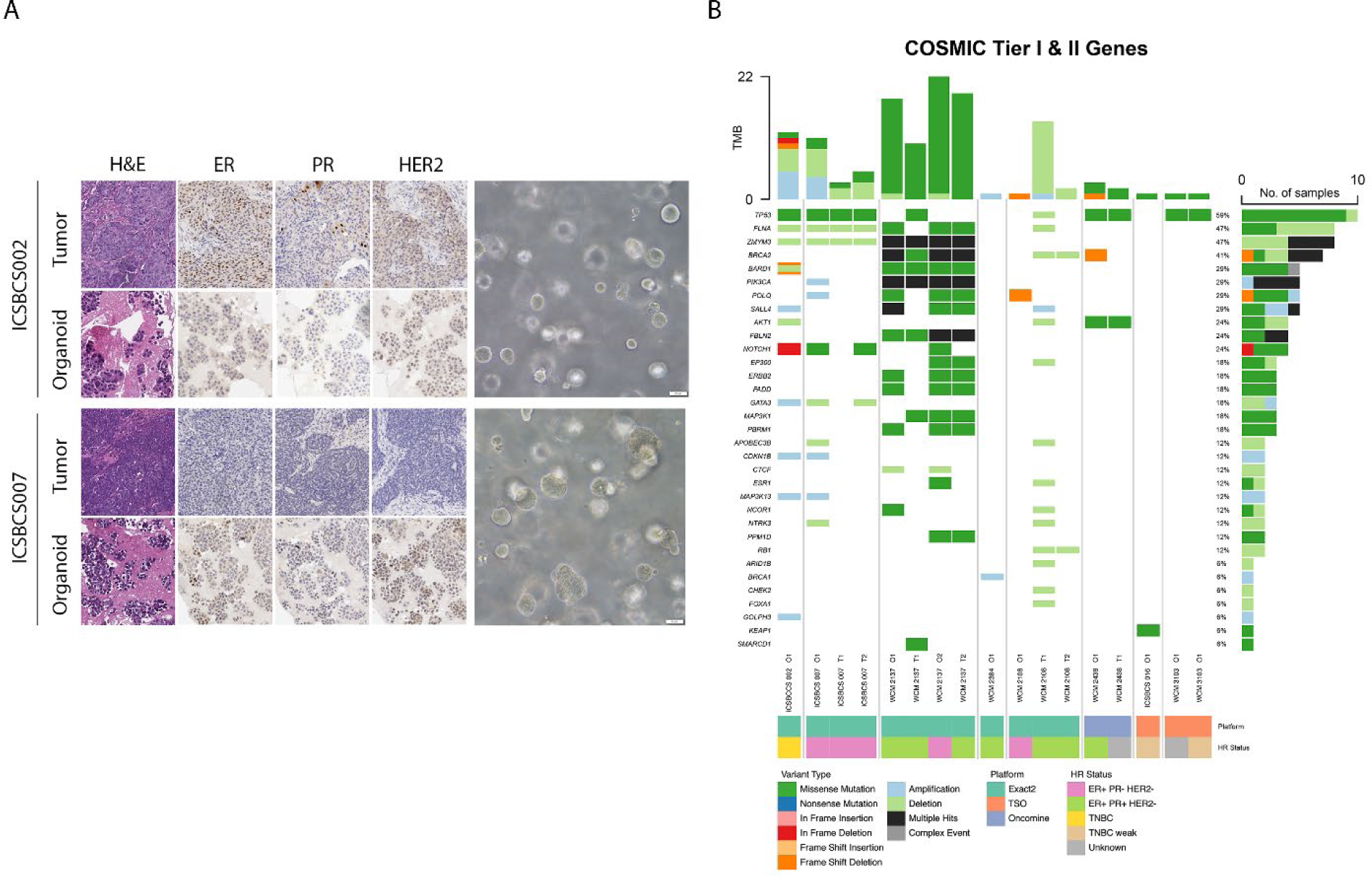
Patient-derived Tumor Organoid characterization. A) Histology and Hormone receptor status of breast PDTO. H&E and immunohistochemical images for breast cancer related markers (ER, PR, and HER2) of tumors and PDTO IHC. Original magnification 20X. B) Oncoplots of COSMIC breast cancer Tier I and II genes. Somatic mutations and copy number alterations from organoids (O) and parent tumor specimens (T) where available (i.e. O1 matches T1, O2 matches T2). Organoids and tumors are columns, and mutated genes are rows. Total tumor mutational burden (TMB) by organoid or tumor specimen is indicated in the top stacked bar chart, where the proportion of specific variant type is indicated. For each row, the total percent mutation across the samples and proportion of variant type is indicated on the right stacked bar chart. Events coded in black indicate multiple hits, where _multiple_ single-nucleotide variant types are detected. Events coded in gray indicate complex events, where both single nucleotide mutations and copy number alterations are detected. Color map column indicates data source for somatic mutation calls as well as molecular subtype of the organoid or parent tumor specimen determined from IHC.

Subtyping was performed by transcriptomic profiling using bulk RNA sequencing to classify the PDTOs into breast cancer intrinsic subtypes based on the PAM50 gene expression panel (**Supplementary Table 2).** PDTOs that were determined to have ER and PR positivity were assigned to be Luminal A (WCM2380, WCM2384), Luminal B (WCM2137_1, WCM2438) or Normal-like (WCM1942, ICSBCS014, WCM2108_2) intrinsic subtypes. PTDOs with ER positivity lacking PR expression were either classified as Luminal (WCM2124,), Luminal B (WCM2137_2) or Basal-like (ICSBCS007, WCM2108_1, with Basal-like having lower *ESR1* gene expression. PDTO ICSBCS002 classified as TNBC by IHC, was determined to be Basal-like by PAM50.

PDTOs from our biobank carried mutations in several breast cancer-relevant genes (**Figure 1B**), including alterations in TP53, BRCA2, and PIK3CA. A comparison of COSMIC tier 1 and 2 mutational profiles between PDTOs and matching patient’s tumors showed 100% concordance for ICSBCS007, WCM2137_2 and WCM2438 PDTOs and 62.5% concordance for WCM2137_1 PDTO (**Figure 1B**, top 50 mutations and additional mutations in **Supplementary Figure 1B,C**). Only one case, WCM2108_1 did not show a good concordance with the tumor. This mutational analysis suggests that the PDTOs are generally representative of the original tumor. We did nonetheless observe that several PDTOs acquired additional mutations compared to the matching tumors (**Figure 1B, Supplementary Figure 1B,C**).

### Kinase genes are differentially expressed in African-ancestry breast cancer patients

Kinases are associated with cancer initiation and progression and several small-molecule kinase inhibitors have been developed with successful application into the clinics. To identify African ancestry-specific associations with kinase genes, we utilized an African-ancestry enriched triple negative breast cancer (TNBC) RNAseq cohort representing West African/Ghanaian (*n* = 6), East African/Ethiopian (*n* = 11), and African American (AA, *n* = 9) patients [28]. Unsupervised hierarchical clustering of 482 kinase genes separated our patients into 3 nodes, where we observed some distinct clustering of Ethiopians, African American and Ghanaians (**Supplementary Figure 2A**). We identified a subset of 80 kinase genes that were significantly associated with self-reported race (SRR) (*p* < 0.05), where unsupervised clustering of the 80 genes more distinctly separate patients into the following nodes with significantly different African (AFR) ancestry levels between the three nodes (*p* = 0.0012) (**Supplementary Figure 2B-C**). A node containing primarily Ethiopian patients (10/11 Ethiopians, 1/9 AA and 1/6 Ghanaians) had lower AFR ancestry (AFR_mean_ = 50.97%). A high AFR node (AFR_mean_ = 96.5%) consisted of primarily Ghanaian patients (4/6 Ghanaians, 1/9 AA). Finally, an intermediate AFR ancestry node (AFR_mean_ = 71.4%) had predominantly AAs (7/9 AA, 1/6 Ethiopians, 1/6 Ghanaians) (**Supplementary Figure 2C**). We next sought to identify therapeutic vulnerabilities in the African ancestry-PDTOs using a domain-focused CRISPR screen, targeting kinase genes.

### A kinome-focused CRISPR-Cas9 screening identifies new vulnerabilities in African-ancestry breast cancer PDTOs

We initially established stable Cas9-expressing clones using a Cas9-P2A-puro lentiviral vector previously described [26]. Due to the extreme sensitivity of breast PDTOs to puromycin selection and subsequent long recovery time; we explored alternative approaches to enrich for Cas9 expression while maintaining the heterogeneity of a polyclonal PDTO population. For this, we linked Cas9 to either: tdTomato or a truncated CD4 protein [31] enabling enrichment of Cas9 expressing cells by flow cytometry or magnetic activated cell sorting (MACS) respectively (**Supplementary Figure 3A**). As expected, Cas9 expression was detected following selection by each approach (**Supplementary Figure 3B**). Both tdTomato and CD4-based Cas9 sorting showed effective depletion of an sgRNA targeting the essential replication factor RPA3 (**Supplementary Figure 3C**), however, isolation of the cells by MACS (CD4-enrichment) was less harsh on cells and allowed rapid enrichment of large numbers of PDTOs. Moreover, this approach enables simple secondary enrichment of Cas9 expressing cells at any point during the screen process in case a sub-population of cells silenced Cas9 (and CD4) expression.

Using truncated CD4 protein as a selection marker, we established stable Cas9 expressing breast PDTOs from ICSBCS002 and ICSBCS007. Cas9 activity was confirmed using reporter genes [32] (**Supplementary Figure 3D**) and by competition assays where we verified that the Cas9 stable lines could induce depletion of sgRNAs targeting a pan-cancer essential gene, RPA3. Cas9 PDTOs that presented at least 80% Cas9 activity based on this assay were selected for screening.

MACS-purified Cas9-expressing PDTOs (ICSBCS002 and ICSBCS007) were transduced with a domain-focused kinome library targeting 482 human kinases (∼6 sgRNAs/gene) [25] at ∼30% infection efficiency with an average representation of 1000 cells/sgRNA (**Figure 2A**). PDTOs were collected 3-4 days post-infection (t0), half of the cells were collected for gDNA, and the rest were expanded for 10 days to allow depletion of sgRNAs targeting pan-essential and organoid-specific essential genes. Cultures were then split across three conditions and treated with either Vehicle (DMSO), the EGFR inhibitor gefitinib, or the MEK1 inhibitor trametinib for 2 weeks. EGFR is frequently over-expressed in TNBC and has been associated with poor prognosis [33] and while EGFR inhibition has been considered a promising approach for TNBC, minimum benefit has been observed in the clinical settings alone or in combination with chemotherapy [34–36]. In addition, EGFR expression has been associated with resistance to hormonal therapy in ER+ breast cancers and has a negative impact on disease-free survival, but EGFR inhibitor treatment in clinical trials showed low clinical benefit rate [37]. Preclinical evidence also supports targeting MAPK cell signaling pathway in TNBC [38] and its inhibition using MEK inhibitors has been studied, but resistance was often observed in pre-clinical studies [39]. This evidence suggests that synergistic combinations with other targeted agents are needed to accomplish clinical efficacy, and we hypothesized that a kinase CRISPR screen in presence of inhibitors of these genes would help uncover strategies for achieving synergy with other kinase inhibitors.

**Figure 2.**
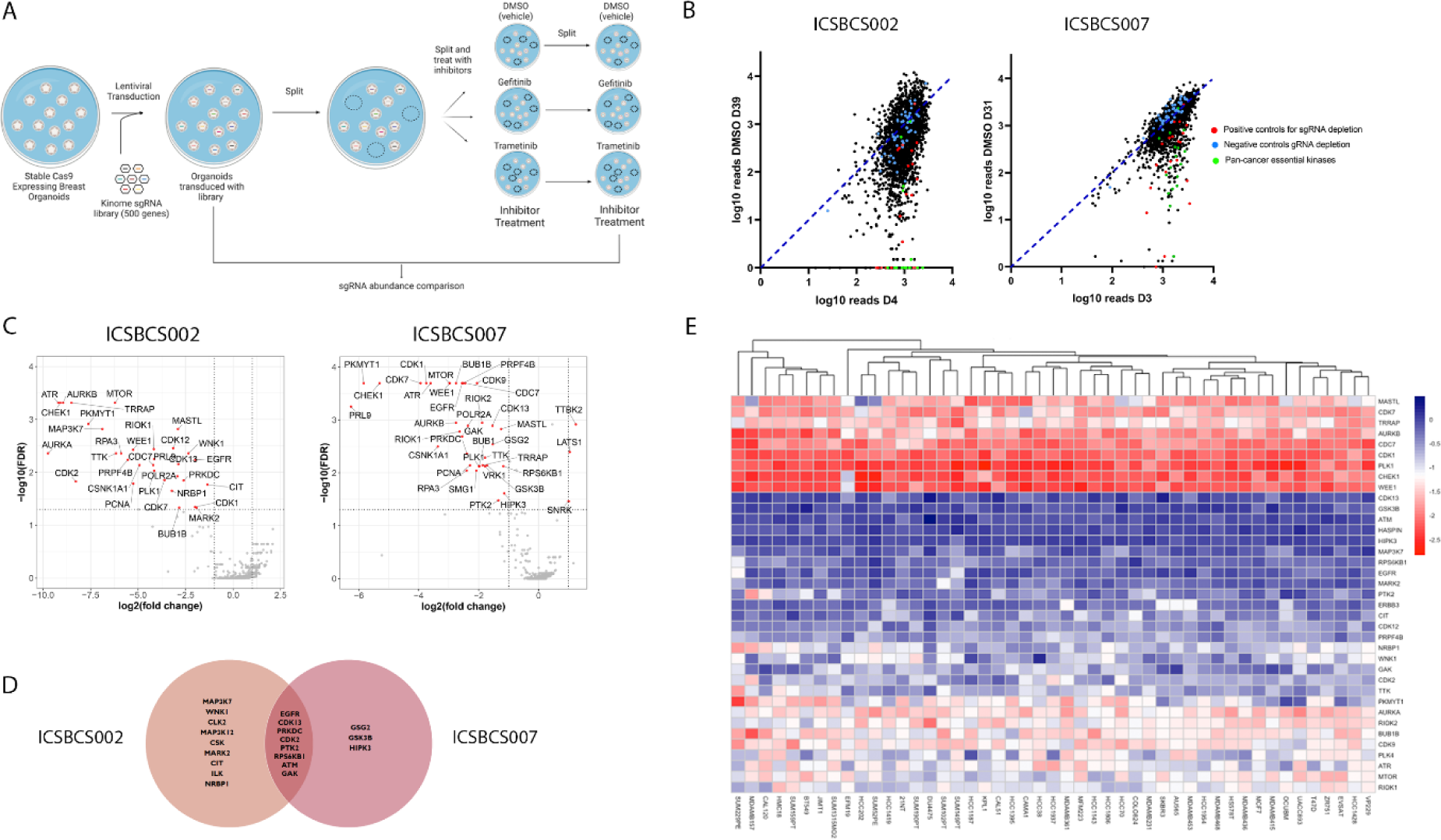
Kinase domain-focused CRISPR-Cas9 Screening in breast cancer organoids. A) Kinase domain-focused CRISPR-Cas9 screening strategy. B) Dot-plot representation of the log10 gRNA counts of DMSO treated PDTO at final time point (D39 or D31) vs initial post-gRNA library transduction (D4 or D3). Indicated are positive (red) and negative (blue) controls for gRNA depletion. gRNAs targeting pan-cancer essential kinases (CHEK1, ATR, AURKA) are depicted in green. C) Volcano plots from the kinome CRISPR screens performed in ICSBCS002 (left) and ICSBCS007 (right). For each gene, the x axis shows its enrichment or depletion, and the y axis shows statistical significance as measured by the false discovery rate (FDR). The horizontal dashed line represents an FDR threshold of 0.05. D) Venn diagrams of non-pancancer essential kinase genes identified in ICSBCS002 and ICSBCS007 (threshold p< 0.05). E) Heatmap showing gene effect for essential hits obtained in CRISPR screening in PDTO in breast cancer cell lines (DepMap data). A lower score means that a gene is more likely to be dependent in each cell line. A score of 0 is equivalent to a gene that is not essential whereas a score of −1 corresponds to the median of all common essential genes.

Drug concentrations used in the screen were determined based on the dose response curves at a value close to the EC_10_ (ICSBCS002: gefitinib 0.5µM, trametinib 6.5 nM and ICSBCS007: gefitinib 0.1µM, trametinib 5 nM) (**Supplementary Figure 4A**). Relative abundance of individual sgRNAs in the DMSO treated samples was compared to initial time point samples (t0) (**Figure 2A, Supplementary table 3**). As expected, sgRNAs targeting known essential genes (positive controls) were strongly depleted, while negative control sgRNAs showed no change in abundance (**Figure 2B**). sgRNAs targeting pan-cancer essential genes that were depleted in 2D cell lines from different tumor types [25, 26], including ATR, CHEK1, AURKA were also depleted in these breast PDTOs (**Figure 2B-C, Supplementary Figure 4B**). In addition, we identified a set of strongly depleted sgRNAs targeting kinases that were not previously identified from 2D cell line studies (**Figure 2C-D, Supplementary Figure 4C**). These putative targets showed similar expression levels between the 2 PDTO lines used, and their expression was stable between passages (passage 5 vs 10) (**Supplementary Figure 4D**). When considering hits as the ones which showed a reduction of at least 50% (log2 fold change <-1) and a p value <0.05, we found several hits shared between the two PDTOs: EGFR, PRKDC, CDK2, PTK2, CDK13, CIT, NRBP1, ILK, RPS6KB1, ATM, GAK (**Figure 2D**). To further confirm the specificity of these depleted gRNAs in regards to breast PDTOs, we compared the hits we obtained in untreated PDTOs (DMSO) in our screens with the gene effect scores for breast cancer cell lines in the Cancer Dependency Map (DepMap) [40, 41] (**Figure 2E**). We found that several of the hits identified in our screen showed minimal effect in DepMap data, perhaps suggesting that either 3D PTDOs and/or cancer models from underrepresented genetic backgrounds may have different genetic dependencies than 2D breast cancer cell lines.

### Inhibitors targeting CDK12/13, CDK2 and ILK validate essential kinase genes

We selected 3 of the putative kinase dependencies identified in both PDTOs to validate the individual gRNAs via a growth assay: CDK2, PTK2 and PRKDC **(Supplementary Figure 4C**). As expected, these individual gRNAs reduced ICSBCS002 PDTO growth, measured as organoid area by high content imaging (**Figure 3A**). We also observed reduced growth in ICSBCS007 PDTO for some of the gRNAs tested for the 3 genes (**Figure 3A-B**).

**Figure 3.**
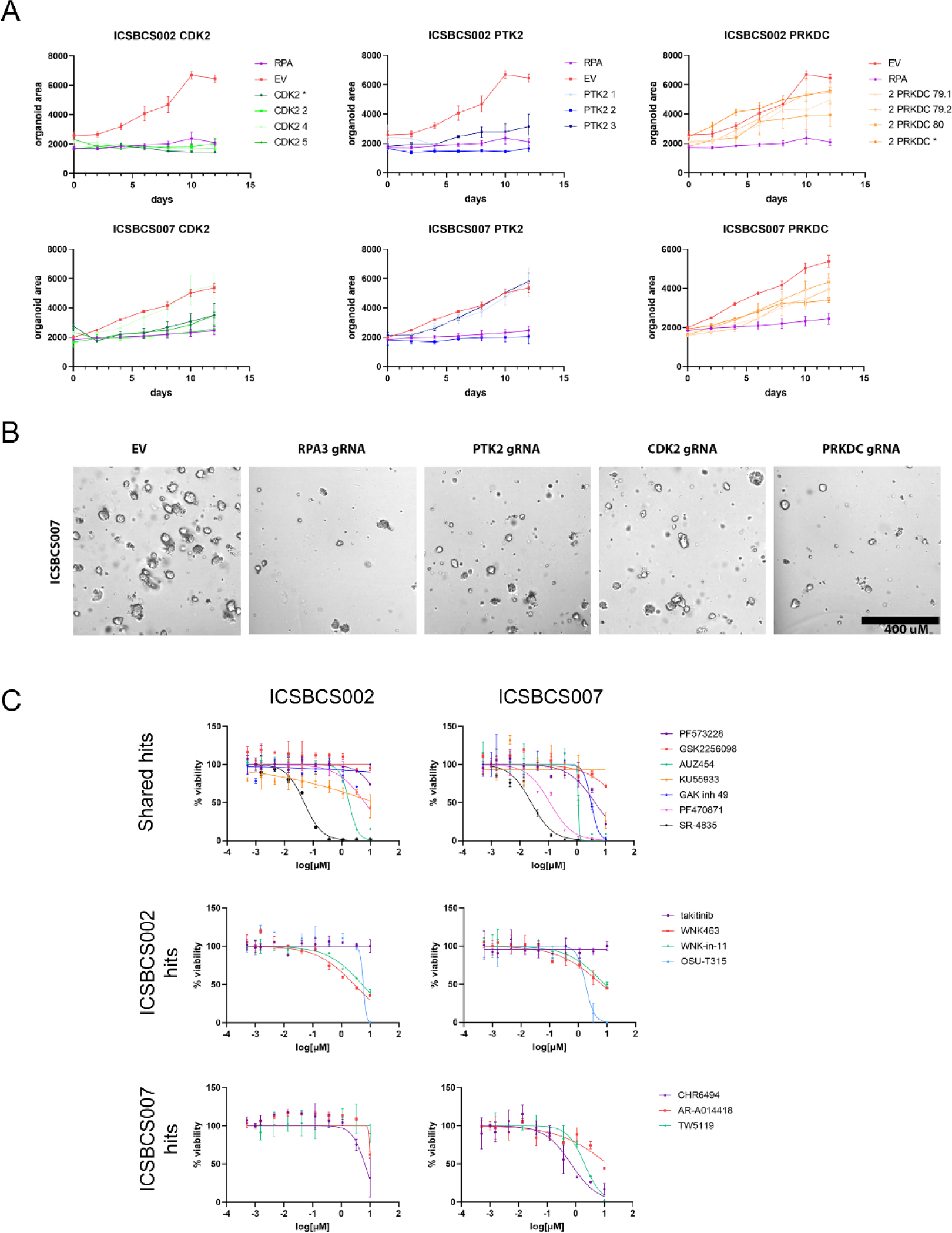
Validation of potential kinase hits in breast cancer organoids and analysis of specific inhibitors. A) Growth assay. ICSBCS002 and ICSBCS007 PDTO were transduced with individual gRNAs against CDK2, PTK2 and PRKDC, empty gRNA control vector (EV) or a positive control with gRNA targeting RPA3 essential gene (RPA) and they were grown and imaged over time to calculate organoid area. B) Representative images of ICSBCS007 PDTO transduced with EV or specific gRNAs. C) Dose response curves to inhibitors targeting potential hits for essential kinases in ICSBCS002 and ICSBCS007. Inhibitors targeting shared hits between the 2 lines (upper panels), ICSBCS002 exclusive hits (middle panel) and ICSBCS007 exclusive hits (lower panel).

To validate pharmacologically a selection of essential kinase hits identified in the 2 breast PDTO lines used in the screens, we tested specific inhibitors against those hits. ICSBCS002 (TNBC) and ICSBCS007 (ER+) were sensitive to several of the tested inhibitors, validating the essentiality of the kinases identified in the screening (**Figure 3C**, IC50 values in S**upplementary Table 4**). Both PDTO lines were sensitive to the CDK2 inhibitor AUZ-454 (**Figure 3C**). We tested 2 different PTK2 (Focal adhesion kinase, FAK) inhibitors, PF-573228 and GSK2256098. Both ICSBCS002 and ICSBCS007 were resistant to GSK2256098, while ICSBCS007 partially responded to PF-573228. The PRKDC specific inhibitors tested (LTURM34 and AZD7648) did not show potent activity against the PDTOs in contrast to the individual gRNA validation results (**Figure 3C**).

Expanding on inhibitors targeting the overlapping hits identified in the CRISPR screens in both PDTO lines, the EGFR inhibitor gefitinib was already included in the screen and both PDTOs were sensitive to it (**Supplementary Figure 4A**). Regarding CDK13, both lines were very sensitive to SR-4835, a CDK12/CDK13 inhibitor (**Figure 3C**). ICSBC007 was sensitive to PF470871, an inhibitor of RPS6KB1 (S6K1) while ICSBCS002 was resistant. ICSBCS002 and ICSBCS007 were resistant to ATM inhibitor KU55933. ICSBCS002 was resistant to GAK inhibitor GAK inhibitor 49, while ICSBCS007 presented some sensitivity (**Figure 3C**).

We tested inhibitors targeting some of the hits identified exclusively for ICSBCS002, MAP3K7 and WNK1. Both ICSBCS002 and ICSBCS007 were resistant to takitinib, a MAP3K7 inhibitor, and to WNK-in-11 and WNK463, WNK1 inhibitors (**Figure 3C**). Both lines presented sensitivity to ILK inhibitor OSU-T315. Even though it was not considered a hit, ILK gRNAs were partially depleted in ICSBCS007 PDTO screen.

We also tested inhibitors targeting certain kinases shown to be essential only in ICSBCS007 PDTOs: GSG-2 and GSK3β. Consistently, ICSBCS007 PDTOs were more sensitive to GSG-2 (Haspin) inhibitor CHR6494 than ICSBCS002 PDTOs and only ICSBCS007 PDTOs presented sensitivity to TWS119 GSK-3β inhibitor (**Figure 3C**). Another GSK3β inhibitor, AR-A014418, was tested but both lines were resistant (**Figure 3C).** These results suggest that the use of inhibitors for EGFR, CDK13, CDK2 and ILK matched the CRISPR screen result for both ICSBCS002 and ICSBCS007. In addition, RPS6KB1, GAK, PTK2, GSG-2 and GSK3β inhibitors effectively inhibit ICSBCS007 PDTOs.

To determine if our candidate kinases are patient-specific novel therapeutic targets, we tested these specific inhibitors in an additional TNBC PDTO line, WCM3103B, and in six ER+ breast PDTOs: WCM2968, WCM2438, WCM2137_2, WCM2108_1, ICSBCS014 and HCM-CSH-0366-C50 (**Figure 4A-B**). WCM2137_2, WCM2108_1 and WCM2968 were derived from African American patients, while ICSBCS014 was obtained from a Ghanaian patient. Like the 2 screened PDTOs, all the other tested breast PDTOs were sensitive to CDK12/13 inhibitor SR-4835 (**Figure 4A-B**, IC50 values in **Supplementary Table 4**), indicating that CDK12/CDK13 represents a potential novel therapeutic target for breast cancer not previously identified in the 2D CRISPR screens reported in DepMap.

**Figure 4.**
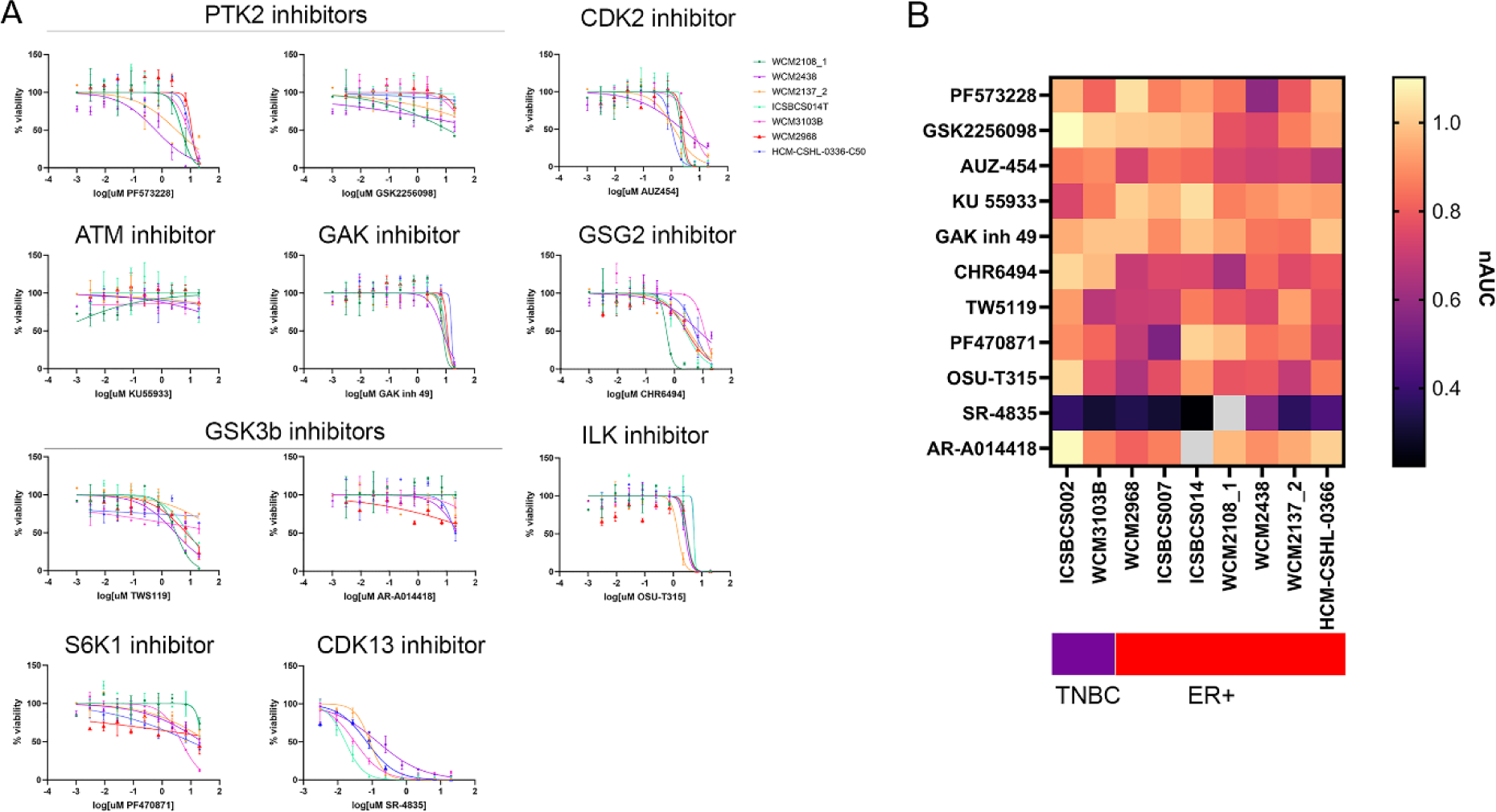
Effect of inhibitors targeting potential essential hits in different breast PDTOs. A) Dose response curves to inhibitors targeting potential essential hits identified in CRISPR screens. TNBC PDTO (WCM3103B) and ER+ PDTO lines (WCM2968, WCM2108_1, WCM2438, WCM2137_2, ICSBCS014, HCM-CSH-0366-C50) were analyzed. B) Heatmap with normalized area under the curve (nAUC) values for the different inhibitors tested in the PDTO lines.

These PDTOs were also sensitive to ILK, CDK2 and GSG-2 inhibitors as ICSBCS002 and ICSBCS007 (**Figure 4A-B**, IC50 values in **Supplementary Table 4**). All the tested lines were sensitive to GAK inhibitor 49, except for ICSBCS002. For PTK2 inhibitors, most of the lines presented sensitivity to PF573228, while all were resistant to GSK2256098. Resistance was also observed in all lines to the ATM inhibitor KU55933. WCM2108_1, WCM2438 and ICSBCS014, responded to GSK-3 inhibitor TWS119 while the rest of the lines did not, and all the lines were resistant to AR-A014418 (**Figure 4A-B**). WCM3103B presented sensitivity to RPS6KB1 inhibitor PF470871 as ICSBCS007, while the other PDTO tested were resistant. These results suggest that some of the essential kinases identified in the CRISPR screening performed in 2 Ghanaian PDTOs, such as ILK, CDK2, GAK, GSG-2, are also essential in other breast PDTO lines, including PDTOs of African ancestry.

### Synergistic interactions between EGFR inhibitor gefitinib and FGFR1

We then sought to identify kinases that were depleted in the context of drug treatment with the EGFR and MEK1 inhibitors, thus suggesting synergistic interactions and opportunities for combination therapies.

In ICSBCS002 PDTO we identified synergy between EGFR inhibitor gefitinib and FGFR1, ERBB2 and Akt2 (**Figure 5A**). In ICSBCS007 PDTO, we found LIMK2 as a hit for synergistic interaction with gefitinib (**Supplementary Figure 5A**). We decided to pursue the validation of the potential synergy between gefitinib and FGFR1, as their synergistic interaction has been previously described in lung cancer cells [42] and several FGFR1 inhibitors are already used in the clinic for the treatment of different malignancies.

**Figure 5.**
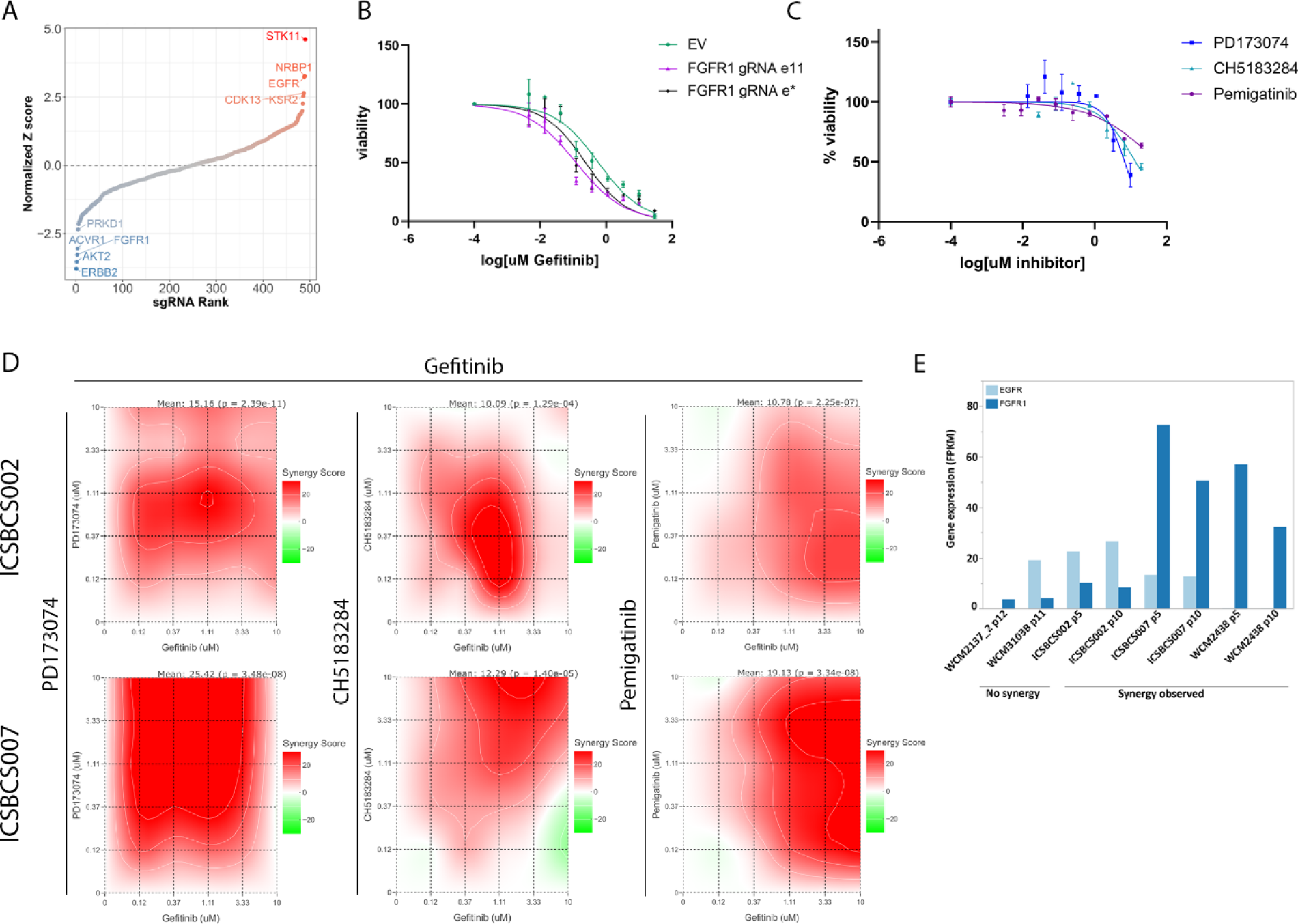
Synergistic interactions with gefitinib. A) Rank plot with normalized Z scores for ICSBCS002 PDTO treated with gefitinib. B) Dose response curves to gefitinib of ICSBCS002 PDTO transduced with individual gRNAs against FGFR1 or empty vector (EV). C) ICSBCS002 drug full dose curves of FGFR1 inhibitors (PD173074, CH5183284 and pemigatinib) and Gefitinib. D) Synergy maps for FGFR1 inhibitors-gefitinib were calculated using the SynergyFinder+ web application with the ZIP synergy model (red indicates a synergistic effect, white an additive effect, and green an antagonistic effect). E) EGFR and FGFR1 gene expression comparison among PDTO samples with and without observed synergy to EGFR and FGFR1 inhibitors. Gene expression of EGFR (light blue) and FGFR1 (dark blue) among PDTO samples. Samples with observed synergy of EGFR and FGFR1 inhibitors contained either EGFR1 amplifications or FGFR1 amplifications.

We performed a CRISPR-Cas9 knock out of FGFR1 in ICSBCS002 PDTO and tested their sensitivity to gefitinib. PDTOs with FGFR1 knock out were more sensitive to gefitinib than cells transduced with an empty vector (IC50 significantly different) (**Figure 5B**) validating the screen results.

To confirm that the synergistic effect of gefitinib in combination with FGFR1 depletion was not solely due to the CRISPR-mediated FGFR1 disruption, we tested the effect of a combination on EGFR and FGFR1 inhibitors. Specifically, we treated ICSBCS002 PDTOs with gefitinib plus different FGFR1 inhibitors: PD173074, zoligratinib (CH5183284) and pemigatinib over a range of concentrations (**Figure 5C** for single drug curves). Utilizing a dose-response matrix, we determined viability using CellTiter-Glo 3D cell viability assay and calculated a synergy score using the SynergyFinder+ web application. We observed synergistic activity following combination treatment of gefitinib with the different FGFR1 inhibitors tested as evidenced by synergy scores greater than 10 (**Figure 5D**).

FGFR1 gene is amplified in ICSBCS007 PDTOs (**Supplementary Figure 1C**). FGFR1 amplification is frequent in breast cancer (10%), predominantly in ER+ breast cancer. It is associated with reduced survival of patients and suggested to have a role in the initiation and progression of breast cancer including *in situ*- to-invasive transition [43, 44]. This amplification may explain why FGFR1 was not identified in the screen. Western blot analysis of ICSBCS007 PDTOs transduced with FGFR1 gRNAs, showed that the gene product was still present, which suggests that the amplified gene was not completely knocked out by the tested gRNAs (**Supplementary Figure 5B**). Despite this, synergy was observed in ICSBCS007 when treated with gefitinib in combination with FGFR1 inhibitors (**Figure 5D**).

We tested if other breast PDTOs also presented synergy between gefitinib and FGFR1 inhibitors. We observed synergy only for a set of concentrations for combinations of gefitinib-PD173074 and gefitinib-CH5183284 in WCM2438 PDTOs (ER+). However, synergy was not observed in 2 other PDTO models tested, WCM3103_B (TNBC, white) and WCM2137_2 (ER+, African American) (**Supplementary Figure 5C**). PDTO lines that presented synergy showed FGFR1 or EGFR amplification (**Supplementary Figure 1C**) and expressed higher levels of those genes (**Figure 5E**).

In the presence of the MEK inhibitor trametinib, we found PRKCG to be a hit in ICSBCS002 PDTOs (**Supplementary Figure 6A**). For ICSBCS007 PDTO, RAF1 was identified as a potential hit for synergistic interaction with trametinib (**Supplementary Figure 6B**). Knock out of RAF1 in ICSBCS007 PDTOs showed a modest increase in sensitivity to trametinib (**Supplementary Figure 6C**). We tested if trametinib presented synergy in combination with a RAF1 specific inhibitor, ZM336372. Synergy for the combination treatment was observed only for a range of concentrations in ICSBCS007 PDTOs (**Supplementary Figure 6D**).

### High-Throughput Drug Screening in the presence of EGFR inhibitor gefitinib

To further investigate potential combinations for EGFR inhibitor gefitinib, we performed a high-throughput drug screen (HTDS) to identify small-molecule compounds that potentiate the response to it in PDTOs. EGFR was identified as a hit in the CRISPR screens in both PDTO lines as a kinase that impacts PDTO viability. Since the EGFR inhibitor gefitinib was already included in the CRISPR screen to identify kinases that may affect sensitivity to it, a HTDS in the presence of gefitinib would allow for the identification of additional synergistic interactions with non-kinase inhibitors as well as to validate its results.

A panel of 156 compounds—including FDA-approved chemotherapeutic and targeted inhibitors and various compounds that target key cancer-related pathways, such as proliferation, apoptosis, DNA repair, and metabolism (listed in **Supplementary Table 5**)—were tested individually or in combination with EGFR inhibitor gefitinib in ICSBCS002 and ICSBCS007. PDTOs were treated for 4 days against a 7-point concentration range of the compound library in combination with a single concentration of gefitinib (IC_30_ concentration: 0.1 µM gefitinib) and cell viability was determined by CellTiter-Glo 3D. To identify combinations that generate synergistic interaction we calculated synergy scores using the highest single agent (HSA) method, to identify combinations that induced higher inhibition than single agents. An average of the HSA score obtained for the range of concentrations tested was plotted (**Figure 6A**).

**Figure 6.**
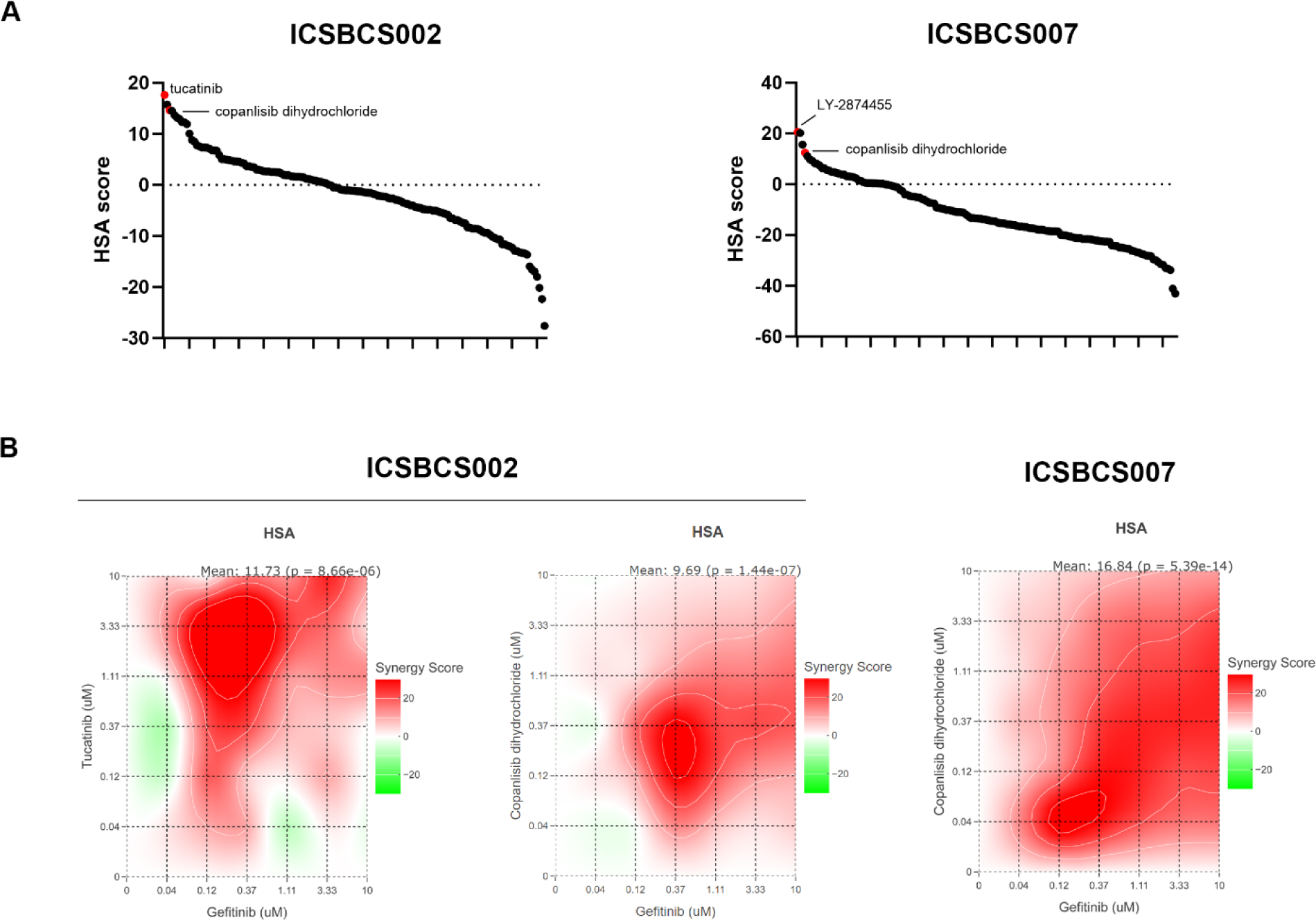
High-throughput drug screening in the presence of EGFR inhibitor gefitinib validates CRISPR screening result and identifies potential synergies. A) The graphs show the average HSA score across the 156 drugs screened in combination with 0.1 µM gefitinib. B) Synergy maps for Tucatinib-gefitinib (left) and Copanlisib dihydrochloride-gefitinib (middle) in ICSBCS002 and Copanlisib dihydrochloride-gefitinib in ICSBCS007 (right) were calculated using the SynergyFinder+ web application with the HSA synergy model (red indicates a synergistic effect, white an additive effect, and green an antagonistic effect).

Tucatinib, an ERBB2 inhibitor, was identified as a sensitizer to gefitinib in ICSBCS002 PDTOs (**Figure 6A, B**). This result validates the kinome CRISPR screening performed in the presence of gefitinib, where ERBB2 was identified in the same PDTO as a hit. Additionally, LY287445, a pan-FGFR inhibitor, was found to potentiate the effect of gefitinib in both lines (**Figure 6A**). This is consistent with the results from the validation of CRISPR screening data, where synergistic interaction was identified between gefitinib and distinct FGFR1 inhibitors. Copanlisib dihydrochloride, a pan-class I PI3K inhibitor, was also identified in the HTDS as synergistic. Overall, these results validate CRISPR screen findings and show additional potential combinations with EGFR and CDK2 inhibitors.

## Discussion

In this study, we developed a breast PDTO biobank focused on an underrepresented group of patients with a focus on African (Ghanaian) and African American patients. A comprehensive characterization of these PDTO models at the genomic, transcriptomic and cytopathological level shows that they faithfully recapitulate the patient tumor fundamental properties. PDTOs are valuable models for precision medicine approaches in drug screening and drug discovery. In addition, this type of initiative also increases visibility of underrepresented patient populations, and it will help to standardize protocols in international collaborations. By using a kinome-focused pooled CRISPR Cas9 screen in a subset of our breast cancer PDTOs, we identified multiple putative therapeutic targets and identified FGFR1 as mechanism of resistance to EGFR inhibition.

Application of CRISPR screening to PTDOs has been challenging due to the 3D configuration of organoids, their heterogeneity that could lead to differences in organoid sizes and growth rates. Moreover, it has been described that when performing CRISPR screens in organoids only a fraction of gRNAs showed functionality in comparison to 2D transformed cell lines [45]. Despite these challenges, data comparing CRISPR screens performed in 3D versus 2D contexts showed that 3D screens can identify vulnerabilities not observed in the 2D cultures [46]. Consistently, by applying CRISPR-Cas9 screening technology in 3D PDTO models from underserved patients, we identified both previously known essential kinases as well as several putative therapeutic targets specific for PDTOs not previously identified in 2D breast cancer lines [41]. The identification of essential kinases not previously identified in 2D breast cancer cell line CRISPR screens, such as EGFR, CDK13, CDK2, GSG2 and S6K1, highlights the importance of establishing physiologically relevant precision medicine pre-clinical models.

Targeted therapies are a core tenet of precision medicine and kinase gene targets have paved the way, with the kinase inhibitor Imatinib being the first targeted therapy developed almost 2 decades ago [47, 48]. Driver mutations in different types of tumors occur in kinase coding genes and there are multiple kinase inhibitors available both in the clinic and under clinical trials [49]. However, in many cases therapeutic resistance is generated to single agent kinase inhibitors by acquisition of mutations or activation of different kinase pathways. In breast cancer, targeted therapies against the kinase gene HER2 are a first-line effective treatment for patients with HER2 dependent tumors [50, 51]. Consistent with our previous study, TNBC RNA-seq indicated that specific kinase genes are differentially expressed between African American and European American patients [29].

There are clear differences in breast cancer incidence and mortality related to race in the United States [52], emphasizing the importance of incorporating diverse populations into molecular studies and clinical trials to address potential biological differences in cancer disparities. Available data from CRISPR screens in cancer cells, such as DepMap project, was generated using cancer cell lines from European or East Asian genetic ancestry, with a very low percentage of African genetic ancestry [53]. Moreover, CRISPR-Cas9 library design should consider germline variants that could likely reduce the cutting activity of Cas9 protein, and this bias affected predominantly cell models derived from African descent individuals as their genomes tend to diverge more from the used consensus genomes [53]. The kinome gRNA library used in our work targets the kinase domain, which we expect to be highly conserved across different races.

The CRISPR screens were performed in 2 breast PDTOs from different breast cancer subtypes, one TNBC and one luminal ER+. Several of the hits identified as dependencies were shared between the two lines and moreover, we did not observe any correlation in sensitivity to the inhibitors tested with the breast cancer subtype. Between the essential kinases identified in both PDTOs, CDK2 and CDK13 were of particular interest as all the tested PDTO lines responded to their respective inhibitors, AUZ-454 and SR-4835 CDK2 has been proposed to have a role in Palbociclib resistance in ER+ breast cancer [54, 55] and in endocrine-therapy resistant breast cancer. CDK2 inhibition has been shown to inhibit proliferation of tamoxifen-resistant cells and restore their sensitivity to anti-estrogen therapy[56]. There are several CDK2 and CDK2/4/6 inhibitors currently being studied in breast cancer resistant to CDK4/6 inhibition [55, 57, 58]. SR-4835, a selective dual inhibitor for CDK12 and CDK13, has been shown to cause TNBC cell death [59] and to act in synergy with DNA-damaging chemotherapy and PARP inhibitors and with anti-PD-1 checkpoint inhibitors [60]. Consistent with our results, CDK12 was previously identified as an essential kinase in different cancer cell lines [26], but DepMap shows that it is not essential for the majority of 2D breast cancer lines. Another kinase identified as essential in both PDTOs was S6K1, which is part of the PI3K pathway signaling pathway and the principal effector of mTORC1. Pharmacologically, we found that ICSBCS007 (ER+) was particularly sensitive to the S6K1 inhibitor PF470871. Different studies suggest that S6K1 has an important role in ER+ breast cancer, as it directly phosphorylates and activates Erα, which promotes S6K1 transcription and mediates cell proliferation [61]. It has also been associated with TNBC, as TNBC cells express high levels of Estrogen related receptor alfa (ERRα) which negatively regulates S6K1 expression. Downregulation of ERRα expression sensitized ERα-negative breast cancer cells to mTORC1/S6K1 inhibitors [62].

For some of the putative kinase hits, we could not replicate the effect of loss of gene function with specific enzymatic inhibitors, for example in the case of PRKDC. One potential explanation for this discrepancy could be that those kinases have a scaffolding role regulating signal transduction by facilitating interaction between different proteins.

Another aim of the CRISPR screen was to identify synergistic interactions between kinases and EGFR and MEK inhibitors. We identified a very promising synergistic interaction between EGFR and FGFR. Synergy between EGFR and FGFR inhibitors has been previously identified in lung cancer, where combined EGFR and FGFR inhibition in FGFR1-overexpressing, EGFR-activated models showed a synergistic effect on tumor growth in cell lines and patient-derived xenografts [42]. In addition, increased FGFR1 expression was detected in EGFR Tyrosine kinase-resistant non-small cell lung cancer (NSCLC) cell lines and it combined EGFR and FGFR inhibition showed some efficacy [63]. Also, EGFR activation has been proposed as an FGFR inhibitor resistance mechanism in gastric cancer [64, 65]. Feedback activation of EGFR signaling was also shown to limit the efficacy of FGFR inhibitor therapy, driving adaptive resistance in patient-derived models of FGFR2 fusion-positive cholangiocarcinoma [66]. In agreement with these studies, our results showed that EGFR and FGFR inhibitors present synergy on breast PDTOs with high expression levels of EGFR and FGFR1. To strengthen the combination potency, it would be necessary to perform the CRISPR screening for longer periods of time, achieving a higher number of population doublings and eliminating false positives.

Here we applied two different strategies for functional precision medicine, a genetic screen and a pharmacological screen. These two approaches showed to be complementary, as different hits were identified but also HTDS in the presence of gefitinib allowed us to validate some of the hits identified in the CRISPR screening, including FGFR1 and ERBB2.

Overall, we showed the feasibility of performing focused CRISPR screening in breast PDTOs. In addition, we were able to identify novel essential hits, not previously identified in 2D lines, which may present clinical relevance. These hits seem to be shared between the TNBC and luminal subtypes (ER+). Further studies including a larger cohort of PDTOs are necessary to determine if these results are specifically related to genetic ancestry.

## Methods

### Patient enrollment

The International Center for the Study of Breast Cancer Subtypes (ICSBCS) is an international consortium of breast cancer clinicians and researchers with the broad goal of studying the heterogeneity of breast cancer across diverse population groups [18, 28, 67]. The ICSBCS obtains written informed consent of patients across all study locations, and all work has been conducted in accordance with established ethical guidelines. Institutional Review Board (IRB) approval has been obtained at all participating study locations, and in the present analysis with patients consented from the United States (Weill Cornell Medical College in New York City, NY) and Ghana (Komfo Anokye Teaching Hospital, Kumasi, Ghana) (IRB #1807019405 and #1305013903).

### PDTO establishment and culture

PDTO lines were developed as previously described [68] with slight modifications. Fresh tissue specimens were collected directly in the procedure rooms. A part of the specimen was immediately frozen in Tissue-Tek OCT Compound (Sakura, 4583), another part was fixed with formalin to generate formalin-fixed paraffin-embedded (FFPE) blocks and the rest of fresh tissue was transported to the laboratory for PDTO establishment after cytopathology review for adequacy [69].

Fresh mastectomy samples were placed in CO_2_ independent medium (Gibco) with GlutaMAX (1×, Invitrogen), 100 U/ml Antibiotic-Antimycotic (Thermo Fisher Scientific) and Primocin 100 μg/ml (InvivoGen). Tissue samples were washed in media two times before being placed in a 10 cm petri dish for mechanical dissection. The dissected tissue was then enzymatically digested with 250 U/ml of collagenase IV (Life Technologies) with 10 μM ROCK inhibitor (Selleck Chemical Inc.) in a 15 ml conical centrifuge tube (Falcon) incubated in a shaker at 37 °C set to 200 rpm. Incubation time of the specimen was dependent on the amount of collected tissue and ranged from 20 to 60 min, until the majority cell clusters were in suspension. After tissue digestion, Advanced DMEM/F12 media (Invitrogen) containing GlutaMAX (1×, Invitrogen), 100 U/ml penicillin-streptomycin (Gibco), and HEPES (10 mM, Gibco) (+++ media) was added to the suspension and the mixture was centrifuged at 300 rcf for 3 min. The pellet was then washed with +++ media and resuspended in breast organoid-specific culture media (**Supplementary Table 6**). The final resuspended pellet was combined with Matrigel (Corning) in a 1:2 volume media:Matrigel, with 5 x 80 μl droplets pipetted onto each well of a six-well suspension culture plate (GBO). The plate was placed into a cell culture incubator at 37 °C and 5% CO_2_ for 30 min to polymerize the droplets before 3ml of breast organoid-specific culture media was added to each well. The culture was maintained with fresh media changed twice a week. Dense cultures with PDTO ranging in size from 200 to 500 um were passaged every 10-30 days depending on the PDTO line. During passaging, the organoid droplets were mixed with TrypLE Express (Gibco) and placed in a water bath at 37 °C for a maximum of 7 min. The resulting cell clusters and single cells were washed and replated, following the protocol listed above. Breast cancer PDTO were biobanked using Recovery Cell Culture Freezing Medium (Gibco) at −80 °C and after 24h transferred to liquid nitrogen. Throughout PDTO development and maintenance, cultures were screened for various Mycoplasma strains using the PCR Mycoplasma detection kit (ABM) and confirmed negative before being used for experimental assays. PDTOs were classified as growing well or long-term cultures if they survived beyond passage 5 (p5), were successfully cultivated through freeze-thaw cycles and showed sustained proliferation without deceleration beyond p5. Short-term cultures are defined as cultures with organoid formation observed but not passaged enough to be confirmed as growing well before taken out of culture.

For PDTO samples collected from Ghanaian patients, we followed the above protocol with the following deviations. After tissue digestion and washing with the +++ media, the single cell suspension was frozen using Bambanker freezing medium (Fujifilm, 302-14681) at −80 °C. Frozen cell suspensions were shipped frozen on dry ice for further processing at WCM. Upon arrival, frozen samples were transferred to −80 °C storage. Single cell suspensions were thawed, washed with +++ media and pellets were resuspended in complete medium and plated as described.

HCM-CSHL-0366-C50 breast PDTO was obtained from ATCC.

### Characterization of PDTOs: Histopathology, DNAseq and RNAseq

#### PDTO Pathological Characterization

PDTO histopathology was verified by comparing sections from fixed passage 5 PDTO cell blocks to parent tumor sections using our developed cytology and histology platforms [70]. For sample preparation, PDTOs were released from Matrigel droplets using Cell Recovery Solution (Corning, Cat# 354253), and pellet was resuspended in a Fibrinogen (Sigma-Aldrich, Cat# F3879)/Thrombin (Sigma-Aldrich, Cat# T4648) (v/v 10:1), and fixed with 4% paraformaldehyde in PBS (Thermo, Cat# J61899AK.) Then, FFPE blocks were made, and stainings were performed. Hematoxylin and eosin (H&E) and Ki67 stained sections of the FFPE blocks were used to verify PDTO as tumor cells and compared to the corresponding tumors to verify matching cellular morphology by a WCM pathologist. Immunohistochemical staining for ER, PR, HER2 was also used for subtyping.

#### Whole exome sequencing and analysis

Whole exome sequencing (WES) was performed using the Exome Cancer Test v2.0 (EXaCT-2) assay that was developed with Agilent based on SureSelect Human All Exon V6 (Agilent Technologies) (manuscript in preparation). EXaCT-2 combines whole exome sequencing with deep sequencing of ∼900 cancer genes. DNA from PDTO, parent tumors and matched normal underwent quality control, where quantification was determined using Qubit assay (ThermoFisher Scientific) and Tapestation 4200 (Agilent Technologies). Samples with a DIN score of <3 were excluded.

Real Time Analysis software (Illumina) was used for primary processing of sequencing images, and samples were demultiplexed and converted to Fastq using bcl2fastq (v2.19, Illumina). Fastp [71] was used to perform quality control analysis and adapter removal, and the processed reads were aligned to the human genome (GRCh37) using bwa mem [72]. Using the GATK toolkit [73] and Picard (http://broadinstitute.github.io/picard). PCR duplicates were marked and removed, and base quality score recalibration was performed using GATK best practices.

After tumor-normal pairs were confirmed from the same individual using SPIA [74], a consensus somatic mutation calling approach was completed as previously described using the TCGA Multi-Center Mutation Calling in Multiple Cancers (MC3) pipeline [75]. Tumor mutational burden (TMB) was determined based on the total number of non-synonymous mutations normalized to the number of bases in exonic regions per million. MSISensor [76] was used to determine microsatellite instability, and copy number variants were determined using CNVKit tools [77]. Copy-number data from parent tumor sample were refined using CLONET tumor ploidy and purity correction [78].

#### Targeted cancer gene panels

Where WES could not be achieved, we employed two targeted clinical genomics assays, the TruSight Oncology 500 (TSO) assay developed by Illumina, and the Oncomine assay developed by ThermoFisher Scientific. DNA extracted from either FFPE or cell pellets was sent to WCM Clinical Genomics Laboratory, where library preparation and TSO and Oncomine sequencing was performed. Both raw data and clinical reports were provided.

#### Somatic variant and CNV visualization

Somatic mutations and copy number alterations from PDTOs and parent tumor specimens where available were obtained from EXaCT-2 WES, or TSO/Oncomine targeted sequencing panels performed by WCM Clinical Genomics Laboratory. Filtered variant and CNV calls obtained from the 3 platforms were visualized as oncoplots using Maftools [79].

#### RNA library preparation and rRNA depletion

RNA was extracted from PDTO samples using Maxwell RSC simply RNA Cells Kit (Promega) and underwent quality control. NanoDrop (ThermoFisher Scientific) was used to measure RNA concentration, and RNA integrity was checked using a 2100 Bioanalyzer (Agilent Technologies). rRNA depletion, RNA sample library and RNA sequencing was completed by the WCM Genomics Core Laboratory. Illumina Ribo Zero Gold for human/mouse/rat kit (Illumina) was used to remove rRNA during sample preparation per manufacturer’s protocol. Library preparation was performed using Illumina TruSeq Stranded mRNA Sample Library Preparation kit (Illumina) following manufacture’s protocol. Normalized cDNA libraries were pooled and sequenced on Illumina NovaSeq 6000 sequencer with pair-end 100 cycles. Raw sequencing reads in BCL format were processed through bcl2fastq (v2.19, Illumina) for Fastq conversion and demultiplexing.

#### RNA sequencing data analysis

RNA sequencing reads were aligned using hg19 reference genome by STAR (v2.7.2b) using STAR-Fusion (v 1.9.1), to generate both alignments and fusion calls. Alignments were sorted and indexed using Samtools (1.14). FPKM was determined after quantification of gene expression alignments using FeatureCounts [80]. To determine gene expression-based subtypes of breast PDTOs, we used the PAM50 intrinsic subtype classifier [81, 82], that assigns tumors into the following 5 subtypes: Luminal A, Luminal B, Her2, Normal-like and Basal-like.

### Drugs

Gefitinib (EGFR inhibitor), trametinib (MEK inhibitor), KU55933 (ATM inhibitor), PF573228 (PTK2 inhibitor), PD173074 and CH5183284 (FGFR1 inhibitors) were purchased from Selleckchem. OSU-T315 (ILK inhibitor), AUZ454 (CDK2 inhibitor), GAK inhibitor 49 (GAK inhibitor), CHR6494 (GSG2 inhibitor), TX1-85-1 (ERBB3 inhibitor), PF4708671 (S6K1 inhibitor), SR-4835 (CDK13 inhibitor), TW5119 and AR-A014418 (GSK3b inhibitors), GSK2256098 (PTK2 inhibitor), pemigatinib (FGFR inhibitor) were purchased from MedChemExpress. Drugs were diluted and utilized according to manufacturers’ instructions. The working concentrations are indicated in each figure panel. Drugs were replenished in the media every 3-4 days if needed.

### Lentiviral transduction

HEK293T cells were plated in T75 flasks at about 90% confluence; 18 hours later cells were transfected using Lipofectamine 3000 (Thermo Fisher Scientific L300008) in the following manner: 10μg lentiviral vector, 7.5μg Pax2, 5μg VSV-G, 30μl of P3000 were combined in 750μl Opti-MEM Medium (Thermo Fisher Scientific 31985070). 30μl Lipofectamine 3000 was diluted in 750μl Opti-MEM, such that the DNA mixture could be diluted 1:1 with the Lipofectamine mixture. Following a 15-minute incubation at room temperature (RT), the transfection mix was added dropwise to the cells while swirling the flask. After 6-7 hours, medium was changed to DMEM (Thermo Fisher Scientific 11965-118) and 10% FBS. Viral supernatants were collected twice every 24h, pooled, concentrated 10x using LentiX concentrator reagent (Takara Bio 631231) and preserved in 0.4 ml aliquots at −80C.

PDTO were transduced with concentrated lentivirus as follows: a single cell suspension was generated, and 500,000 cells were plated per well of a 48 well suspension plate and the viral supernatant dilution was added with 12μg/mL polybrene (EMD Millipore TR-1003-G). The plate was centrifuged at 600 g for 40 minutes at 32C followed by a 3-4 h incubation at 37C. Cells were then collected in Eppendorf tubes and washed with +++ media, resuspended in breast media-Matrigel and plated in 60ul droplets in a 12-well plate.

### Generation of Cas9-expressing clones

Breast PDTO cells were transduced with lentiviral vector pLenti-Cas9-P2A-ΔCD4 and selected using magnetic associated cell sorting (MACS) with anti-CD4 antibodies conjugated with nanobeads (Miltenyi). As part of the optimization process, PDTO were transduced with plenti-Cas9-P2A-puro vector [83] and were selected with 1 µg/ml puromycin or plenti-p2A-tdtomato and selected by Fluorescence-activated cell sorting. PDTOs were expanded and tested for Cas9 activity utilizing control gRNA (empty vector, EV) or an gRNA targeting RPA3 (positive control, essential gene). PDTO that showed good depletion of RPA as compared to the control were expanded and utilized for the pooled negative-selection human kinome gRNA-based screen.

### Pooled negative-selection human kinome gRNA based screen

A human kinase domain-focused gRNA library (Addgene 117725) [25] was used to assess CRISPR Cas9 screening as a tool in breast cancer PDTO to identify whether novel vulnerabilities could be detected for each PDTO. The CRISPR library pool was transduced into stable Cas9 expressing PDTO lines at infection rates between 30-50% to ensure single lentiviral integration per cell and a coverage of 1000 cells per gRNA (1000x). 3 days after lentiviral transduction, PDTO lines were split, and cells were sampled via flow cytometry to determine frequency of fluorescently linked guides. Half of the cells (6 million transduced cells) were snap-frozen and kept for sequencing as original library representation (Day 3-4), and the rest was allowed to recover for 7 days to ensure crRNA-based genome editing. At 10 days (D10) post library transduction the cells were replated in the presence of DMSO, gefitinib, and trametinib. Drugs were replenished after 4 days. After another 14 days (D31-D39) in the presence of DMSO and/or the inhibitors, cells were snap frozen for the final time points (D31-DMSO, D31-GEF, and D31-TRM). Genomic DNA was extracted from Day2, Day31-DMSO, Day31-GEF and Day31-TRM using QIAGEN DNA mini kit. 55 parallel PCRs were performed using 5.5ug of total gDNA as a template, pooled using PCR purification kit (QIAGEN) and eluted in 85ul PCR-grade water. After template repair, dA tailing, barcoding and pre-capture PCR as described [26], libraries were quality-checked using Bioanalyzer. Sequencing was performed by the WCM Genomics Core Laboratory. ICSBCS002 samples were sequenced with NextSeq500 mid output (150-bp single-end reads) and ICSBCS007 samples with NextSeq500 high output (150-bp single-end reads). Throughout the screen, organoid media was changed every 3-4 days.

The sgRNA counts were analyzed with the MAGeCK v0.5.9.4 [84] and the DrugZ v1.1.0.2 [85] algorithms. Synergistic interactions were identified by pairwise comparison of treatment and vehicle groups with MAGeCK Robust Rank Aggregation (RRA). Datasets were also analyzed by DrugZ to similarly identify drug synergy. Genetic perturbations with false discovery rates (FDR) of <0.05 were considered potential hits. Gene essentialities were identified with MAGeCK RRA pairwise comparison of vehicle treated samples at indicated time point to either day 3 or day 4 samples.

### Cloning

gRNAs targeting genes for validation were cloned into the LRT2B lentiviral backbone using BsmBI and adapters as described previously [86]. All vectors and primers used are listed in **Supplementary Table 7 and 8**.

### Drug dose-response assay (manual)

PDTO were digested into a single cell suspension and cells were plated in a 384 well plate at a density of 1200 cells per well in 8ul droplets (1:2 media:matrigel). Plates were centrifuged briefly to ensure that the cells were at the bottom of the well and 15μl of media were added. Cells were incubated for 72h to allow PDTO formation and after the incubation the drugs were added. Plates were incubated for an additional 96h. After incubation the readout was performed using CellTiterGlo®3D reagent according to the manufacturer’s protocol. Luminescence was measured by the Biotek Synergy H4 plate reader. Non-linear regression analysis, IC50 determination and area under the curve calculation were performed using GraphPad Prism 9 software. We defined the PDTOs as sensitive to an inhibitor when viability was reduced below 50%.

For synergy analysis, the Synergyfinder+ web application [87] was used. The synergy score was calculated by using the zero interaction potency (ZIP) Synergy model. A score lower than −10 indicates an antagonistic effect (green area), between −10 and +10 indicates an additive effect (white area), and higher than +10 indicates a synergistic effect (red area).

### High-throughput drug screening (HTDS)

HTDS was run using a specially equipped HighRes Biosolutions automation platform controlled by Cellario software. Along with integrating Hamilton Microlab Star liquid handler, this robotic system included Prime Automated liquid handler, HighRes Biosolutions Microspin plate centrifuge, plate incubators, Biotek Synergy H4 plate reader and high content imager (Operetta CLS, Perkin Elmer). PDTOs were seeded at predetermined density based on optimal proliferation (1200/well) for each cell line in 384-well plates by Prime Automated Liquid Handler and the drugs were added using automated liquid handling (Hamilton) after 72h. A library of 156-FDA approved drugs and various compounds that target key cancer-related pathways (**Supplementary table 5**) was tested as single agents and in combination with Gefitinib at IC30 concentration (0.1uM) over a 7-point concentration range. PDTOs were incubated 96h in the presence of drugs. Viability was assessed with CellTiterGlo®3D. Quality control was performed based on the coefficient of variation (CV) of the DMSO control cells (CV <3%). Highest single agent (HSA) score was calculated by using Synergyfinder plus web application 3.0 [87].

### Viability and growth assay

A single cell suspension of cells transduced with the indicated gRNAs or EV were plated in a 384 well plate at a density of 1200 cells per well in droplets of 8μl containing 66.6% Matrigel with 30μl of media for 2 weeks changing media twice a week. PDTO were imaged every 2-3 days using a high content imager (Operetta). Bright field images were analyzed by the Harmony software to determine organoid area.

### Flow cytometry

For the Fluorescence-based competition assay, lentiviral supernatants were titrated such that PDTOs expressing Cas9 would achieve a 30-60% tdTomato+ or GFP+ cells at day 3 post-transduction. The day 3 timepoint was used for normalization for all subsequent time points such that D3 represents 100%. Cells were split at D3, D17, and D32, 1/5th of the cells were replated at each time point and the remainder were analyzed as single cells via flow cytometry on a BD LSRFortessa Flow Cytometer.

For the fluorescence-based Cas9 activity assay, PDTO were transduced with lentiviral supernatants for pKLV2-U6-gRNA5(Empty)-PGK-BFP-2A-GFP-W and pKLV2-U6-gRNA5(gGFP)-PGK-BFP-2A-GFP-W (Addgene plasmids # 67979 and# 67980) [32]. After 3 to 5 days post lentiviral transduction, cells were analyzed for GFP and BFP expression via flow cytometry on a BD LSRFortessa Flow Cytometer.

### Western blot

Cells from the indicated conditions were harvested from 1 well of a 6 well suspension plate. PDTOs were dissociated with TrypLE (Gibco 12605-028), in the manner described above, and resuspended in 150ul RIPA buffer, incubated 15min on ice and centrifuged at 13.000rpm to collect protein lysates. 30ug per lane were loaded; the following antibodies were used: anti-FGFR1 (Cell Signaling Technology 9740S), anti-Cas9 (BioLegend 844301), goat anti-rabbit HRP (Abcam ab205718), goat anti-mouse HRP (Thermofisher Scientific 32230), and β-Actin monoclonal HRP (Thermo Fisher Scientific MAF-15739-HRP).

## Acknowledgements

This work was supported by the Englander Institute for Precision Medicine. Project support for this research was also provided in part by the Center for Translational Pathology from the Department of Pathology and Laboratory Medicine at Weill Cornell Medicine.

