## Supplementary Figures for "Kinome focused CRISPR-Cas9 screens in African ancestry patient-derived breast cancer organoids identifies essential kinases and synergy of EGFR and FGFR1 inhibition"

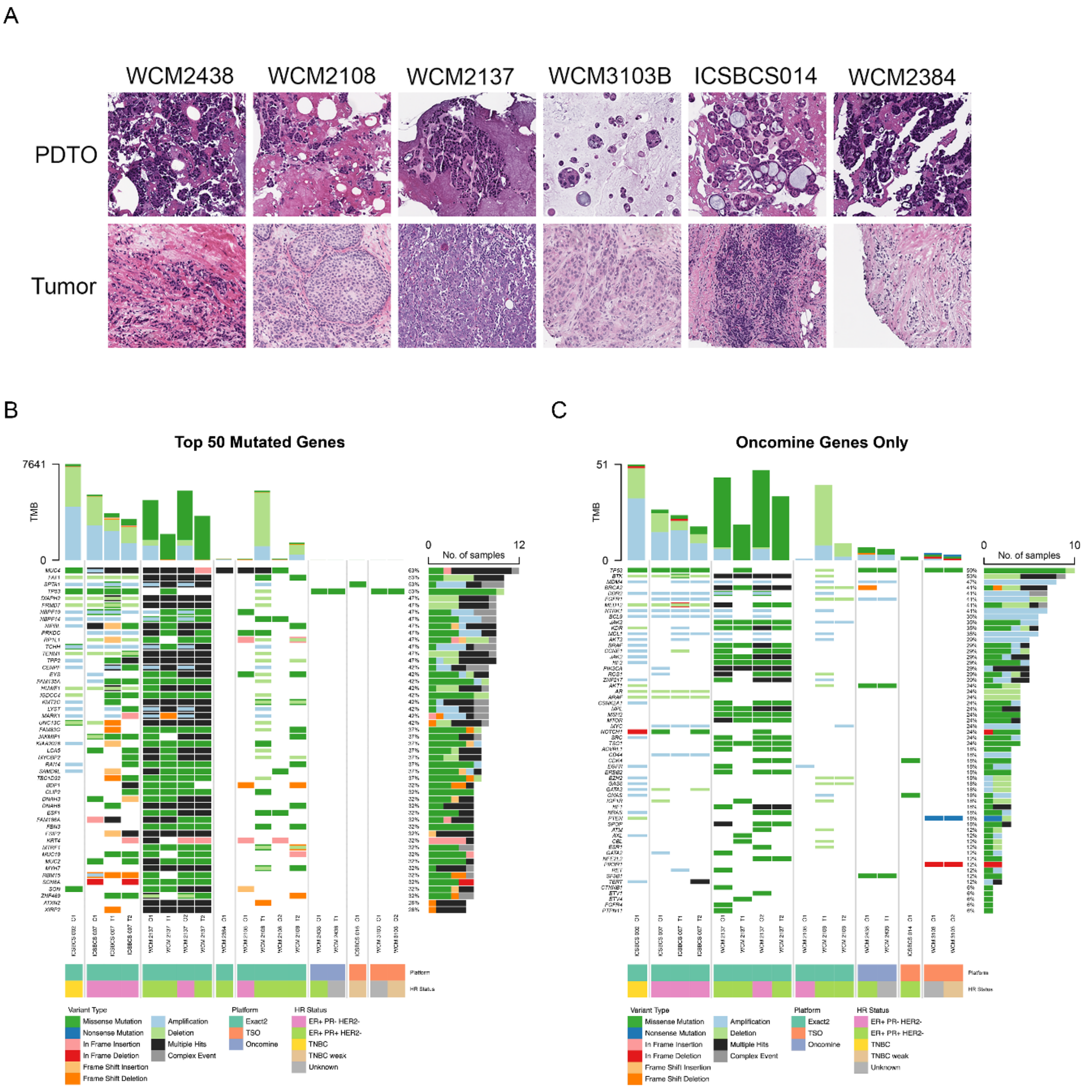


**Supplementary Figure 1**. **PDTOs characterization**. A) Histological images of hematoxilin & eosin staining of breast cancer PDTOs and matching tumor. B) Mutational profile of PDTO and matching tumors. Oncoplots of Top 50 mutated genes, and C) Oncomine targeted gene panel, including clinically relevant genes. Somatic mutations and copy number alterations from organoids (O) and parent tumor specimens (T) where available as described in Figure 1B.

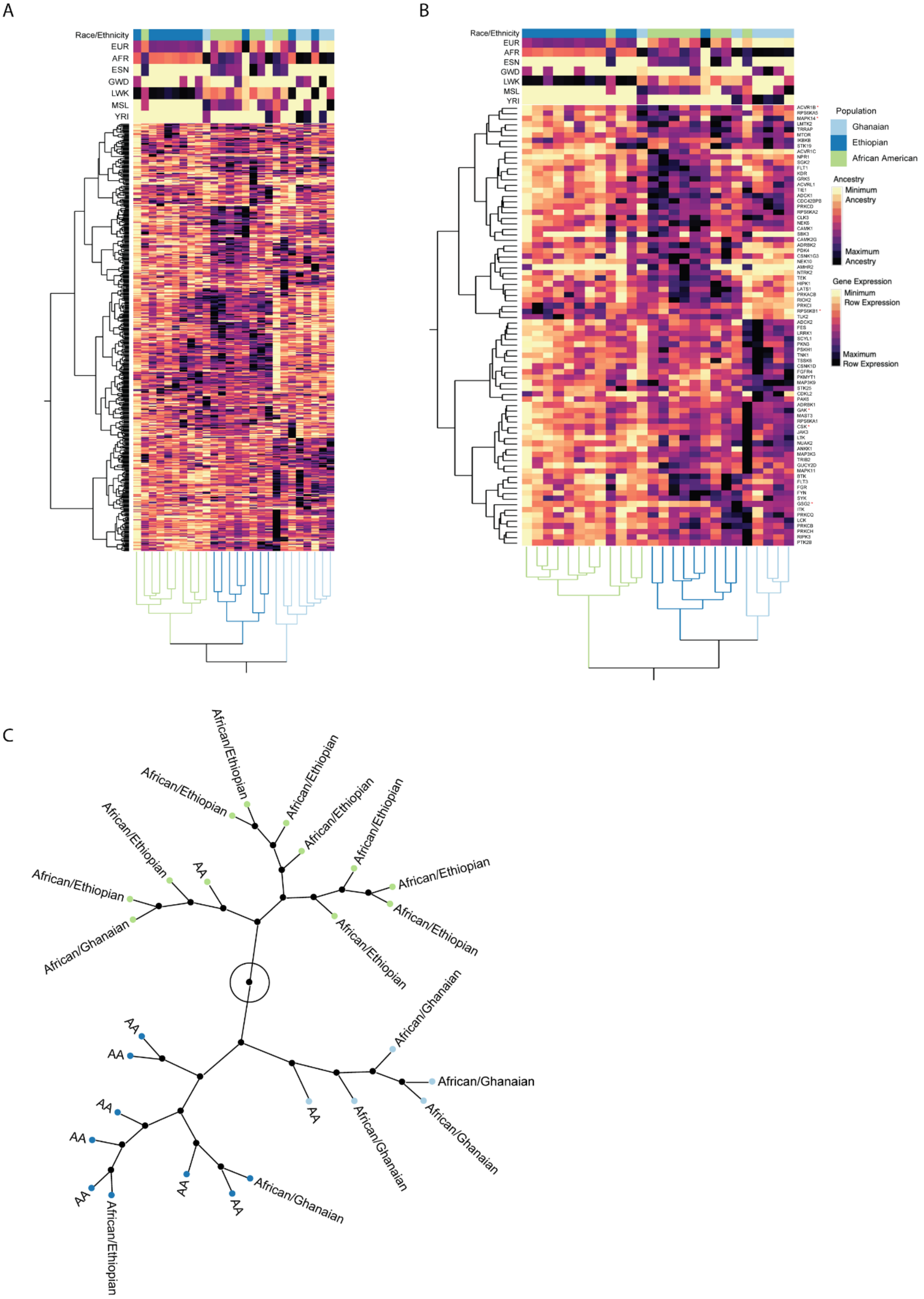

A
B
Race/Ethnicity
EUR
AFR
ESN
GWD
LWK
MSL
YRI
Population
Ghanaian
Ethiopian
African American
Ancestry
Minimum Ancestry
Maximum Ancestry
Gene Expression
Minimum Row Expression
Maximum Row Expression
C
African/Ethiopian
African/Ghanaian
AA

**Supplementary Figure 2. Kinase expression in an African ancestry enriched TNBC cohort.** A) Unsupervised hierarchical clustering of 482 kinase genes included in CRISPR PDTO screen in an African ancestry enriched TNBC cohort. Patients are columns, and genes are rows. Color map indicates patient self-reported race, quantified genetic ancestry. (B) Unsupervised hierarchical clustering of 80 SRR-associated kinase genes included in CRISPR PDTO screen in an African ancestry enriched TNBC cohort. Patients are columns, and genes are rows. Color map indicates patient self-reported race, quantified genetic ancestry. Genes marked with a red star were also identified as essential MAGeCK essential genes. (C) Constellation plot based on clustering of 80 significantly associated kinase genes. Three nodes shown in (B) displayed as a constellation plot. Color of dots corresponds to color of patient clusters.

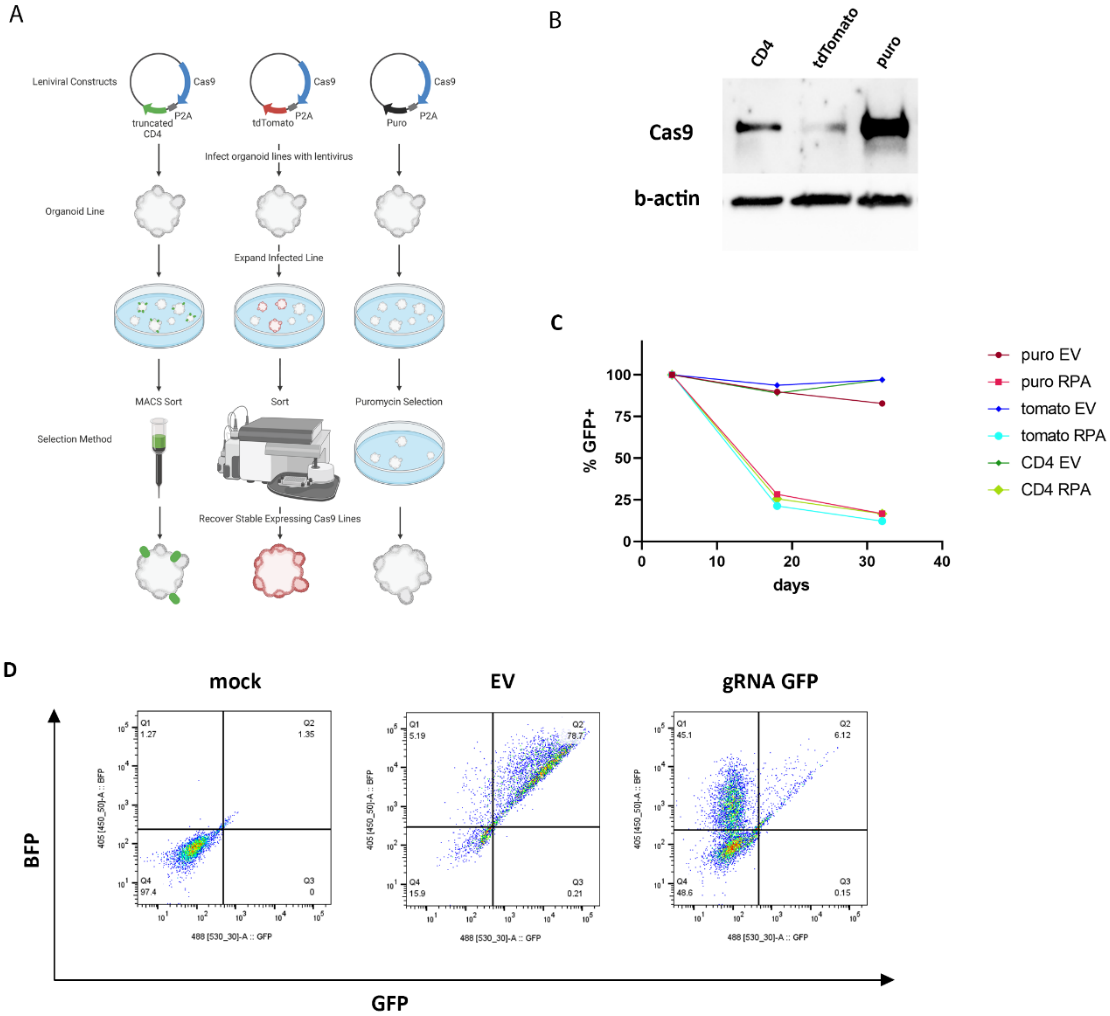


**Supplementary Figure 3**. **Generation of Cas9-stable breast PDTO lines.** A) Strategies for generating stable Cas9 organoid lines. B) Western Blot showing Cas9 expression in ICSBCS002 organoids expressing Cas9-ΔCD4, Cas9-tdtomato and Cas9-puro vectors. C) Fluorescent competition assay to measure Cas9 activity in ICSBCS002 organoids. Cells expressing RPA3 sgRNAs were outcompeted by EV-transduced cells over 2-3 passages (around 30 days in culture, depending on the organoid line), as shown by flow cytometry-tracking of GFP expression. EV= empty gRNA control vector, RPA= RPA3 gRNA vector. D) Validation of Cas9 activity in ICSBCS007 organoids using the Cas9 activity reporter. Representative dot plots.

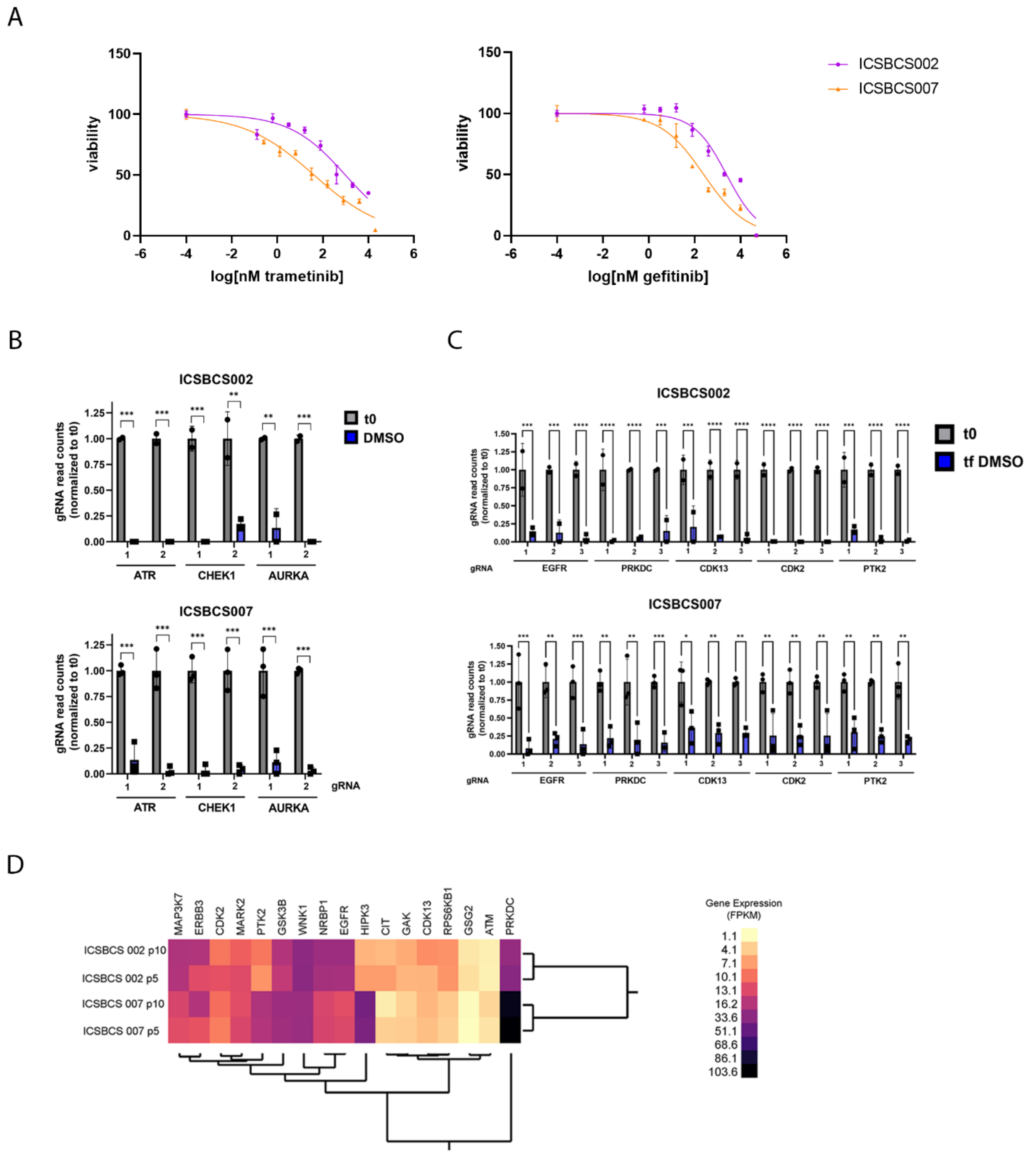


**Supplementary Figure 4. Kinome-focused CRISPR screening in PDTO.** A) Dose response curves to trametinib and gefitinib. B) Normalized gRNA counts at initial time point (t0, gray) and DMSO treated (tf DMSO, blue) breast cancer organoids for ATR, CHEK1 and AURKA. Results are shown normalized to initial time point, n= 2 or 3 independent experiments, as mean and SEM. C) Normalized gRNA counts at initial time point (t0, gray) and DMSO treated (tf DMSO, blue) breast cancer organoids for EGFR, PRKDC, CDK13, CDK2 and PTK2. Results are shown normalized to initial time point, n= 2 or 3 independent experiments, as mean and SEM.D) Expression of MAGeCK novel essential genes identified in Ghanaian PDTO breast models. Unsupervised hierarchical clustering of MAGeCK essential genes in the ICSBCS 002 and ICSBCS 007 PDTO models at an FDR $p < 0.05$ threshold. Expression data for passage 5 and passage 10 is shown (p5 and p10, respectively). Gene expression is in FPKM.

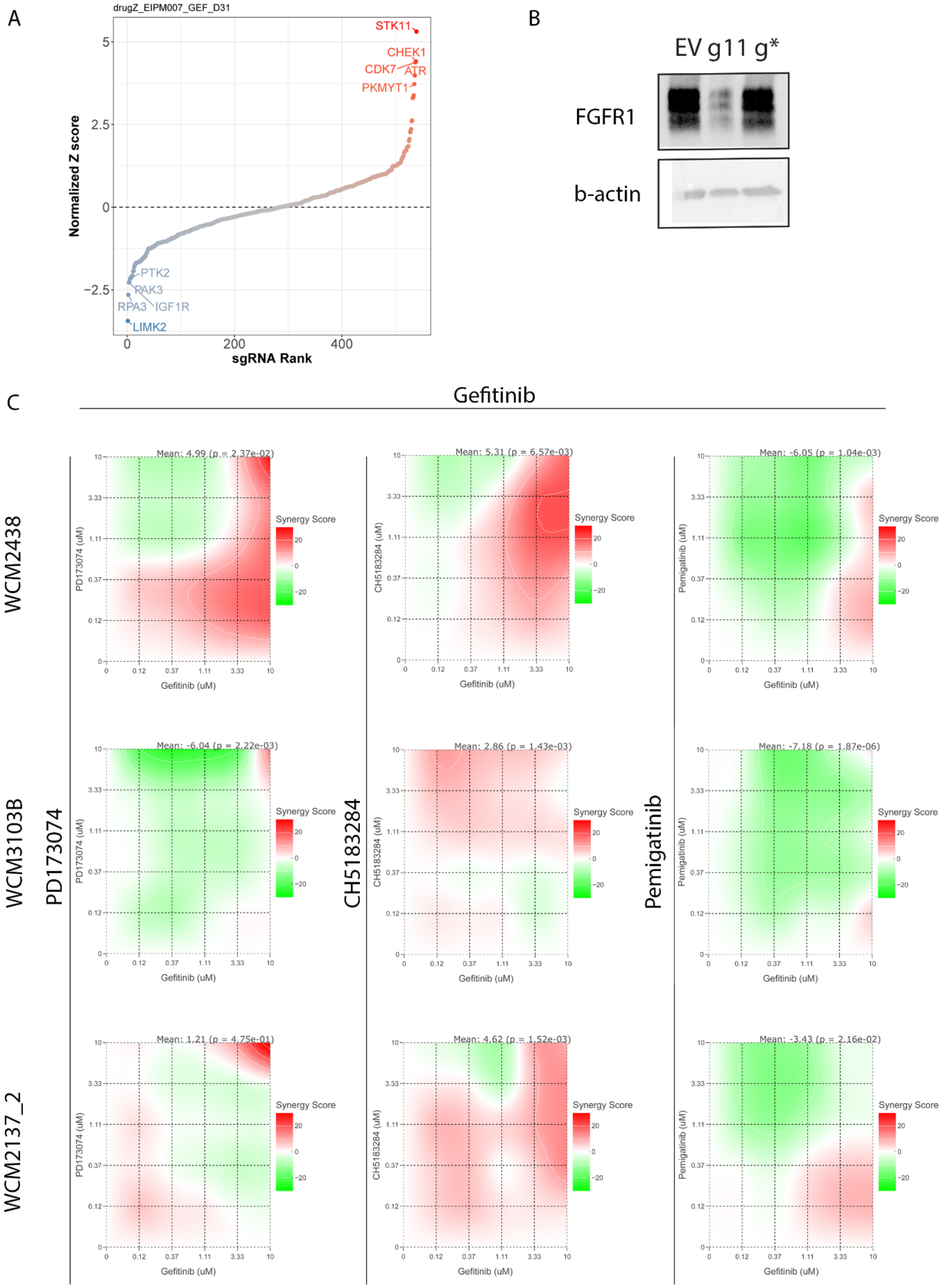

A
drugZ_EIPM007_GEF_D31
Normalized Z score
sgRNA Rank
STK11
CHEK1
CDK7
ATR
PKMYT1
PTK2
PAK3
RPA3
IGF1R
LIMK2
B
EV g11 g*
FGFR1
b-actin
C
Gefitinib
WCM2438
WCM3103B
WCM2137_2
PD173074
CH5183284
Pemigatinib
Mean: 4.99 (p = 2.37e-02)
Mean: 5.31 (p = 6.57e-03)
Mean: -6.05 (p = 1.04e-03)
Mean: -6.04 (p = 2.22e-03)
Mean: 2.86 (p = 1.43e-03)
Mean: -7.18 (p = 1.87e-06)
Mean: 1.21 (p = 4.75e-01)
Mean: 4.62 (p = 1.52e-03)
Mean: -3.43 (p = 2.16e-02)
Synergy Score
PD173074 (uM)
CH5183284 (uM)
Pemigatinib (uM)
Gefitinib (uM)

**Supplementary Figure 5. CRISPR screen in the presence of gefitinib.** A) Rank plot with normalized Z scores for ICSBCS007 PDTO treated with trametinib. B) Western blot showing FGFR1 expression in ICSBCS007 PDTO transduced with empty vector (EV) or gRNAs targeting FGFR1 (E11 and E*). C) Synergy maps for FGFR1 inhibitors-gefitinib in WCM2438, WCM3103B and WCM2137_2 PDTO lines. Synergy scores were calculated using the SynergyFinder+ web application with the ZIP synergy model (red indicates a synergistic effect, white an additive effect, and green an antagonistic effect).

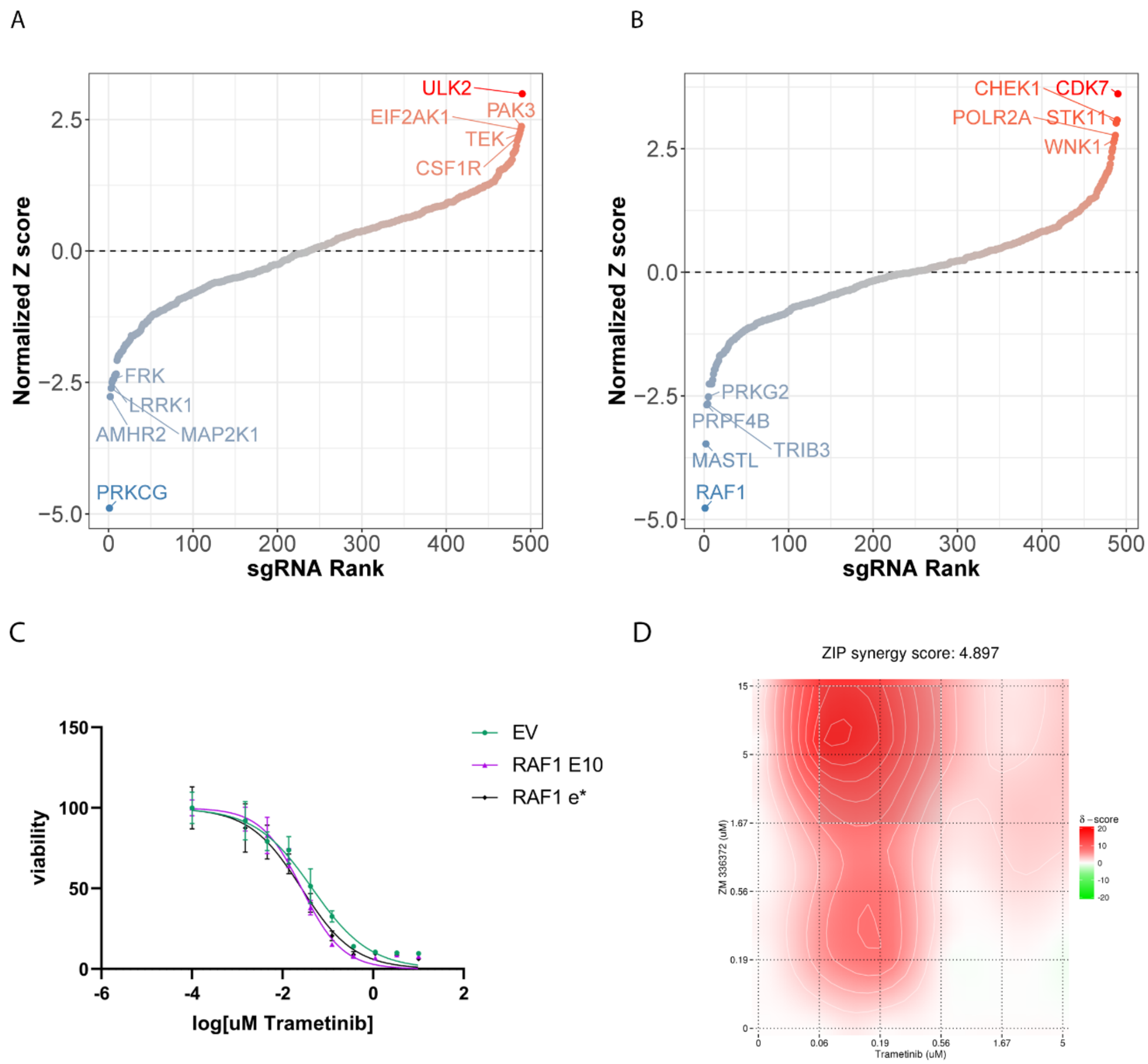


**Supplementary Figure 6. Screening results for synergy in combination with trametinib.** A) Rank plot with normalized Z scores for ICSBCS002 PDTO treated with trametinib. B) Rank plot with normalized Z scores for ICSBCS007 PDTO treated with trametinib. C) Dose response curves to trametinib of ICSBCS007 PDTO transduced with individual gRNAs against RAF1 or empty vector (EV). D) Synergy maps for ZM336372 RAF1 inhibitor-trametinib were calculated using the SynergyFinder web application with the ZIP synergy model (red indicates a synergistic effect, white an additive effect, and green an antagonistic effect).
