## Supplementary Tables 1,2,4,6,7,8 for "Kinome focused CRISPR-Cas9 screens in African ancestry patient-derived breast cancer organoids identifies essential kinases and synergy of EGFR and FGFR1 inhibition"

**Supplementary Table 1. Patient clinical information**

| Patient ID | Race | Ethnicity | Gender | Age | Tumor type | Receptor status tumor | Sample collected pre/post treatment? | Has this patient been diagnosed with metastatic disease? |
|---|---|---|---|---|---|---|---|---|
| ICSBCS002 | Black | Non-Hispanic | Female | unknown | Invasive Ductal Carcinoma | ER+,PR+,HER2- | unknown | unknown |
| ICSBCS007 | Black | Non-Hispanic | Female | unknown | Invasive Ductal Carcinoma | ER+,PR-,HER2- | unknown | unknown |
| ICSBCS014 | Black | Non-Hispanic | Female | 42 | Invasive Ductal Carcinoma | ER-, PR-, HER2- | unknown | unknown |
| WCM1942 | White | Non-Hispanic | Female | 29 | Ductal Carcinoma | ER+, PR+, HER2- | pre | No |
| WCM2108 | Black | Non-Hispanic | Female | 46 | Invasive Ductal Carcinoma | ER+, PR+, HER2- | pre | No |
| WCM2124 | White | Non-Hispanic | Female | 23 | Invasive Ductal Carcinoma | ER+, PR+, HER2+ | post | Yes |
| WCM2137 | Black | Non-Hispanic | Female | 44 | Invasive Ductal Carcinoma | ER+, PR+, HER2- | post | No |
| WCM2384 | Other | Hispanic or Latino | Female | 64 | Invasive Ductal Carcinoma | ER+, PR+, HER2 equivocal | pre | Yes |
| WCM2438 | Other | Declined | Female | 53 | Ductal carcinoma | ER+,PR+,HER2- | unknown | unknown |
| WCM2968 | black | Non-Hispanic | Female | 59 | Invasive Ductal Carcinoma | ER-, PR-, HER2- | post | unknown |
| WCM3103 | white | Non-Hispanic | Female | 80 | Invasive Ductal Carcinoma | ER-, PR-, HER2- | pre | unknown |
| WCM2380 | black | Non-Hispanic | Female | 78 | Carcinoma | ER+, PR+, HER2- | pre | No |

**Supplementary Table 2. PDTOs breast cancer subtype analysis**

| PDTO | Passage | Tumor HR Status | Organoid HR Status | Organoid Ki67 Status | PAM50 Call |
|---|---|---|---|---|---|
| ICSBCS002 | p5 | ER+, PR+, HER2- | ER-, PR-, HER2- | 95% | Basal |
| ICSBCS007 | p5 | ER+, PR-, HER2- | ER+, PR-, HER2- | 95% | Basal |
| ICSBCS014T | p7 | ER-, PR-, HER2- | ER+, PR+, HER2- | 20% | Normal |
| WCM1942 | p5 | ER+, PR+, HER2- | ER+, PR+, HER2- | 40% | Normal |
| WCM2108_1 | p6 | ER+, PR+, HER2- | ER+, PR-, HER2- | 30% | Basal |
| WCM2108_2 | p5 | ER+, PR+, HER2- | ER+, PR+, HER2 equivocal | 35% | Normal |
| WCM2124 | p5 | ER+, PR+, HER2+ | ER+, PR-, HER2- | 20% | LumA |
| WCM2137_2 | p12 | ER+, PR+, HER2- | ER+,PR-,HER2- | | LumB |
| WCM2137_1 | p5 | ER+, PR+, HER2- | ER+, PR+, HER2 equivocal | 50% | LumB |
| WCM2380 | p7 | ER+, PR+, HER2- | ER+, PR+, HER2- | 15% | LumA |
| WCM2384 | p10 | ER+, PR+, HER2 equivocal | ER+, PR+, HER2- | 40% | LumA |
| WCM2438 | p5 | | ER+, PR+, HER2- | 90% | LumB |
| WCM3103B | p11 | ER-, PR-, HER2- | ERlow, PR- ,HER2- | 75% | Her2 |

**Supplementary Table 3. List of raw gRNA counts** (see additional file)

**Supplementary Table 4. IC50 values obtained for tested inhibitors in breast PDTO lines.**

| Inhibitor | WCM2108_1 | WCM2438 | ICSBCS002 | ICSBCS007 | WCM2137_2 | ICSBCS014 | WCM3103B |
|---|---|---|---|---|---|---|---|
| PF573228 | 5.02 | 0.5973 | 23.21 | 4.245 | 3.218 | 4.783 | 8.443 |
| AUZ-454 | 2.149 | 2.396 | 1.818 | 1.117 | 1.265 | 2.207 | 5.666 |
| GAK inhibitor 49 | 8.208 | 7.974 | >100 | 3.12 | 9.27 | 10.17 | 10.54 |
| CHR6494 | 0.5243 | 7.176 | 6.545 | 0.6887 | 3.154 | 2.414 | 11.42 |
| TW5119 | 3.107 | 2.925 | Unstable | 2.03 | 106 | 8.338 | 137 |
| PF470871 | 27.49 | 24.84 | 6.469 | 0.1117 | 40.29 | 23.53 | 4.707 |
| OSU-T315 | 2.989 | 2.584 | 5.733 | 1.791 | 1.454 | 5.35 | 3.039 |
| SRR-4835 | | 0.1013 | 0.04869 | 0.02498 | 0.0857 | 0.0167 | 0.03137 |

**Supplementary Table 5. List of inhibitors tested in HTDS alone or in combination with gefitinib.** (see additional file)

**Supplementary Table 6**. **PDTO breast media composition.**

| Media component | | Final concentration |
|---|---|---|
| Advanced DMEM/F12 | Thermo Fisher Scientific 12634028 | |
| Glutamax | Life Technologies 35050079 | 1% |
| HEPES | Thermo Fisher Scientific 15630-080 | 1% |
| Penicillin/Streptomycin | Thermo Fisher Scientific 15140163 | 100U/ml |
| B27 supplement | Life Technologies 17504-044 | 1X |
| Nicotinamide | Sigma-Aldrich N0636-100G | 10mM |
| N-Acety-l-cysteine | Sigma-Aldrich A9165-5G | 1.25 mM |
| Primocin | Invivogen ant-pm-1 | 1X |
| recombinant human FGF-basic | Peprotech 100-18B | 1 ng/mL |
| recombinant human FGF10 | Peprotech 100-26 | 20ng/ml |
| PGE2 | R&D Systems 2296/10 | 1µM |
| SB202190 | Sigma-Aldrich S7067 | 10µM |
| mouse recombinant EGF | Thermo Fisher Scientific PMG8043 | 50ng/mL |
| Y-27632 | VWR S1049-50MG | 10µM |
| A-83-01 | VWR 10188-672 | 500nM |
| Heregulin beta-1 | PeproTech 100-03 | 200ng/ml |
| Noggin conditioned media | | 10% |
| R-spondin conditioned media. | | 5% |

**Supplementary Table 7**. **List of primers used in this study.**

| Primer | Use | Sequence |
|---|---|---|
| RPA3e1.1_F | gRNA cloning into LRT2B | caccgCCGGCGTTGATGCGCGACCT |
| RPA3e1.1_R | gRNA cloning into LRT2B | aaacAGGTCGCGCATCAACGCCGGc |
| CDK2*_F | gRNA cloning into LRT2B | CACCGGTGCAGAAATTCAAAAACC |
| CDK2*_R | gRNA cloning into LRT2B | AAACGGTTTTTGAATTTCTGCACC |
| CDK2_2_F | gRNA cloning into LRT2B | CACCGATCTCTCGGATGGCAGTAC |
| CDK2_2_R | gRNA cloning into LRT2B | AAACGTACTGCCATCCGAGAGATC |
| CDK2_4_F | gRNA cloning into LRT2B | CACCGTCATGGGTGTAAGTACGAAC |
| CDK2_4_R | gRNA cloning into LRT2B | AAACGTTCGTACTTACACCCATGAC |
| CDK2_5_F | gRNA cloning into LRT2B | CACCGTTTCAGGAGCTCGGTACCAC |
| CDK2_5_R | gRNA cloning into LRT2B | AAACGTGGTACCGAGCTCCTGAAAC |
| PRKDC_79.1_F | gRNA cloning into LRT2B | CACCGCCCAAGCGCATCATCATCCG |
| PRKDC_79.1_R | gRNA cloning into LRT2B | AAACCGGATGATGATGCGCTTGGGC |
| PRKDC_79.2_F | gRNA cloning into LRT2B | CACCGTCATCCGTGGCCATGACGAG |
| PRKDC_79.2_R | gRNA cloning into LRT2B | AAACCTCGTCATGGCCACGGATGAC |
| PRKDC_80_F | gRNA cloning into LRT2B | CACCGTAAGCCGCCTTCTCCTCTT |
| PRKDC_80_R | gRNA cloning into LRT2B | AAACAAGAGGAGAAGGCGGCTTAC |
| PRKDCe*_F | gRNA cloning into LRT2B | CACCGTTTTAGAAGTTCTAGACTCA |
| PRKDCe*_R | gRNA cloning into LRT2B | AAACTGAGTCTAGAACTTCTAAAAC |
| PTK2_1_F | gRNA cloning into LRT2B | caccgTTACCTCAGCTAGTGACGTA |
| PTK2_1_R | gRNA cloning into LRT2B | aaacTACGTCACTAGCTGAGGTAAc |
| PTK2_2_F | gRNA cloning into LRT2B | caccgTACTAAGCTGATAGGCATAC |
| PTK2_2_R | gRNA cloning into LRT2B | aaacGTATGCCTATCAGCTTAGTAc |
| PTK2_3_F | gRNA cloning into LRT2B | caccgCTTTGGATTATCCCGATATA |
| PTK2_3_R | gRNA cloning into LRT2B | aaacTATATCGGGATAATCCAAAGc |
| FGFR1e*_F | gRNA cloning into LRT2B | CACCGTCAGGGTCAGTTTGAAAAGG |
| FGFR1e*_R | gRNA cloning into LRT2B | AAACCCTTTTCAAACTGACCCTGAC |
| FGFR1 E11 F | gRNA cloning into LRT2B | CACCGCCACTTTGGTCACACGGTT |
| FGFR1 E11 R | gRNA cloning into LRT2B | AAACAACCGTGTGACCAAAGTGGC |
| LRG_F2 | PCR-out sgRNAs from library Rxn | TCTTGTGGAAAGGACGAAACACCG |
| LRG_R2 | PCR-out sgRNAs from library Rxn | TCTACTATTCTTTCCCCTGCACTGT |

**Supplementary Table 8. List of vectors used in this study.**

| Vector | Type | Reference |
|---|---|---|
| pLenti-Cas9-P2A-Puro (Cas9-Puro) | Lentiviral constitutive Cas9 expression vector | Addgene no. 110839 |
| pLenti-Cas9-P2A-tdTomato (Cas9-tdTomato) | Lentiviral constitutive cas9 expression vector | Generated through Vector Builder |
| pLenti-Cas9-P2A-tCD4 (Cas9-tCD4) | Lentiviral constitutive Cas9 expression vector | Generated through Vector Builder |
| pLenti-U6-tdTomato-P2A-BlasR (LRT2B) | Lentiviral gRNA Expression Vector with Tomato and Blasticidin | Addgene no. 110854 |
| LRG 2.1 | Lentiviral gRNA expression vector with GFP | Addgene no. 108098 |
| pKLV2-U6-gRNA5(Empty)-PGK-BFP-2A-GFP-W (Empty-BFP-GFP) | Lentiviral empty gRNA vector control for GFP/BFP Cas9 activity assay | Addgene no. 67979 |
| pKLV2-U6-gRNA5(sgGFP)-PGK-BFP-2A-GFP-W (gGFP-BFP-GFP | Lentiviral sgGFP expressing vector for GFP/BFP cas9 activity assay | Addgene no. 67980 |
